# Flexible, Scalable, High Channel Count Stereo-Electrode for Recording in the Human Brain

**DOI:** 10.1101/2022.11.08.515705

**Authors:** Keundong Lee, Angelique C. Paulk, Yun Goo Ro, Daniel R. Cleary, Karen J. Tonsfeldt, Yoav Kfir, John Pezaris, Youngbin Tchoe, Jihwan Lee, Andrew M. Bourhis, Ritwik Vatsyayan, Joel R. Martin, Samantha M. Russman, Jimmy C. Yang, Amy Baohan, R. Mark Richardson, Ziv M. Williams, Shelley I. Fried, Hoi Sang U, Ahmed M. Raslan, Sharona Ben-Haim, Eric Halgren, Sydney S. Cash, Shadi. A. Dayeh

## Abstract

Over the past decade, stereotactically placed electrodes have become the gold standard for deep brain recording and stimulation for a wide variety of neurological and psychiatric diseases. Current electrodes, however, are limited in their spatial resolution and ability to record from small populations of neurons, let alone individual neurons. Here, we report on a novel, reconfigurable, monolithically integrated human-grade flexible depth electrode capable of recording from up to 128 channels and able to record at a depth of 10 cm in brain tissue. This thin, stylet-guided depth electrode is capable of recording local field potentials and single unit neuronal activity (action potentials), validated across species. This device represents a major new advance in manufacturing and design approaches which extends the capabilities of a mainstay technology in clinical neurology.

**One-Sentence Summary:** A human-grade thin-film depth electrode offers new opportunities in spatial and temporal resolution for recording brain activity.

## Main Text

Brain disorders severely interfere with quality of life and can lead to major socioeconomic disparities (*1, 2*). A major therapeutic approach for a wide variety of neuropsychiatric diseases involves invasive recordings from both the cortex and subcortical structures and/or direct electrical neuromodulation of those structures. For treating medically intractable epilepsy, for example, it is commonplace for recordings to be made using stereotactically placed electrodes (sEEG or depth electrodes). Similarly, electrodes of this type are used to target the thalamus, substantia nigra and other subcortical structures for the control of seizures, Parkinson’s disease, and essential tremor as well as a growing number of other disorders (*3-13*). Future applications of these electrodes could be to understand memory disorders and assist in memory restoration (*14-16*) while other uses could be the development of brain computer interfaces to restore movement and communication in the setting of trauma, amyotrophic lateral sclerosis and stroke (*17*). Electrodes of this type are implanted through small openings in the skull and penetrate the brain parenchyma at varying depths depending on the surgical target, and allow for subcortical recordings and, sulcal depth evaluation, with deep structural reach that is not attainable by surface electrodes. Currently, arrays of electrodes are hand-assembled 0.8-1.27 mm diameter cylinders comprised of 8-16 contacts each 3-5 mm in length.

The manufacture of clinical electrode arrays has only incrementally advanced since their initial development in the early 1950’s because of the limitations in hand assembly and wiring of these implantable devices. In addition, the construction of these electrodes limits their spatial resolution; they are only able to record local field potentials (LFPs) over relatively large areas (e.g. multiple mm) and are unable to record from small, discrete neuronal populations let alone individual neurons (e.g. action potential activity). A variety of modifications of this electrode have been used to record highly local sites in the brain. For example, platinum-iridium microwires extruding from the tip of depth electrodes enable recording of single and multi-unit activity from up to 9 microwires (*18*). This configuration only allows recording from the tip. Dixi Medical has produced a depth electrode with extensible microwires from the body of the array (*19*). Neither of these approaches allows more than a few channels to be recorded, neither afford grid-like high spatial resolution in that developing a spatial map of multiple action potential sites of origin are not possible, and the devices are still hand-made. Other electrodes that can record single units from the human brain and afford high resolution spatial mapping of single cell activity include the Utah array (*20*) and Neuropixels (*21, 22*) with up to hundreds of channels (*23, 24*). These devices, currently used in research, are limited by the silicon (Si) manufacturing technology and the brittleness of Si. They are also currently only able to access superficial cortical layers of the brain.

To increase the spatial resolution and channel count of electrodes that can record from either the lateral grey matter or deep brain structures, recent engineering approaches have focused on rolling or adhering conformable and photolithographically defined polyimide electrodes around or on medical-grade tubing used in clinical depth electrodes (*25-28*). These hybrid integration approaches impose a limitation on the size of the electrode such that the starting diameter is pre-determined by the clinical depth electrode diameter.

To address these various limitations and go well beyond current capabilities, we developed an entirely new manufacturing method for thin-film electrodes enabling reproducible, customizable, and high throughput production of electrodes (1) to be implanted in the operating room using similar brain implant techniques to standard clinical depth electrodes, and (2) to reach deep brain structures and achieve high spatial resolution and channel count with a much thinner electrode body. This new manufacturing process exploits (1) titanium (Ti) sacrificial layers employed in the microfabrication of free-standing microelectromechanical systems (MEMS) devices. A stylet inserted where the Ti sacrificial layer is removed assists in hardening and implanting the depth electrode – similar to the standard clinical SEEG electrode implantation procedures – and is subsequently removed. (2) This MEMS process is implemented on relatively large (18 × 18 cm^2^) glass substrates of (**Fig. 1A**) allowing us to produce multiple copies of the SEEG devices using materials that are typical for manufacturing of display screens. Therefore, this new manufacturing method of thin-film based and clinical-grade depth microelectrode array, termed a micro-stereo-electro-encephalography (μSEEG) electrode, enables flexibility in design, scalability afforded by the display screen manufacturing which is cost effective, and does not involve manual assembly typical for standard SEEG electrodes. The μSEEG dimensions can be made custom for application-based contact spacing and channel count. Here, we illustrate the flexibility of our design by manufacturing and testing μSEEG electrodes ranging from a few millimeters to tens of centimeters long, 1.2 mm wide, and 15 μm thick. The manufacturing is compatible with novel electrode materials that can be used to produce microscale electrode contacts with low electrochemical impedance. We demonstrate the μSEEGs with two low-impedance contact materials, (1) the platinum nanorod (PtNR) contact technology (**Fig. 1B**) we developed (*29, 30*) and (2) the poly(3,4-ethylenedioxythiophene) polystyrene sulfonate (PEDOT:PSS) electrode technology (*31-34*) to record broadband neuronal activity including single units (action potentials) and LFPs in rats, pigs, non-human primates (NHPs), and humans. We also test and demonstrate the flexibility of the manufacturing process which can involve either polyimide or parylene C as the device substrate, both of which are biocompatible. This newly integrated μSEEG electrode induced less tissue damage than cylindrical clinical electrodes in a 2-week rat implant (n = 1). Such a flexible, high channel count system paves the way for expanded and more efficacious neuronal recordings and neuromodulation across the spectrum of neuropsychiatric diseases.

**Fig. 1.**
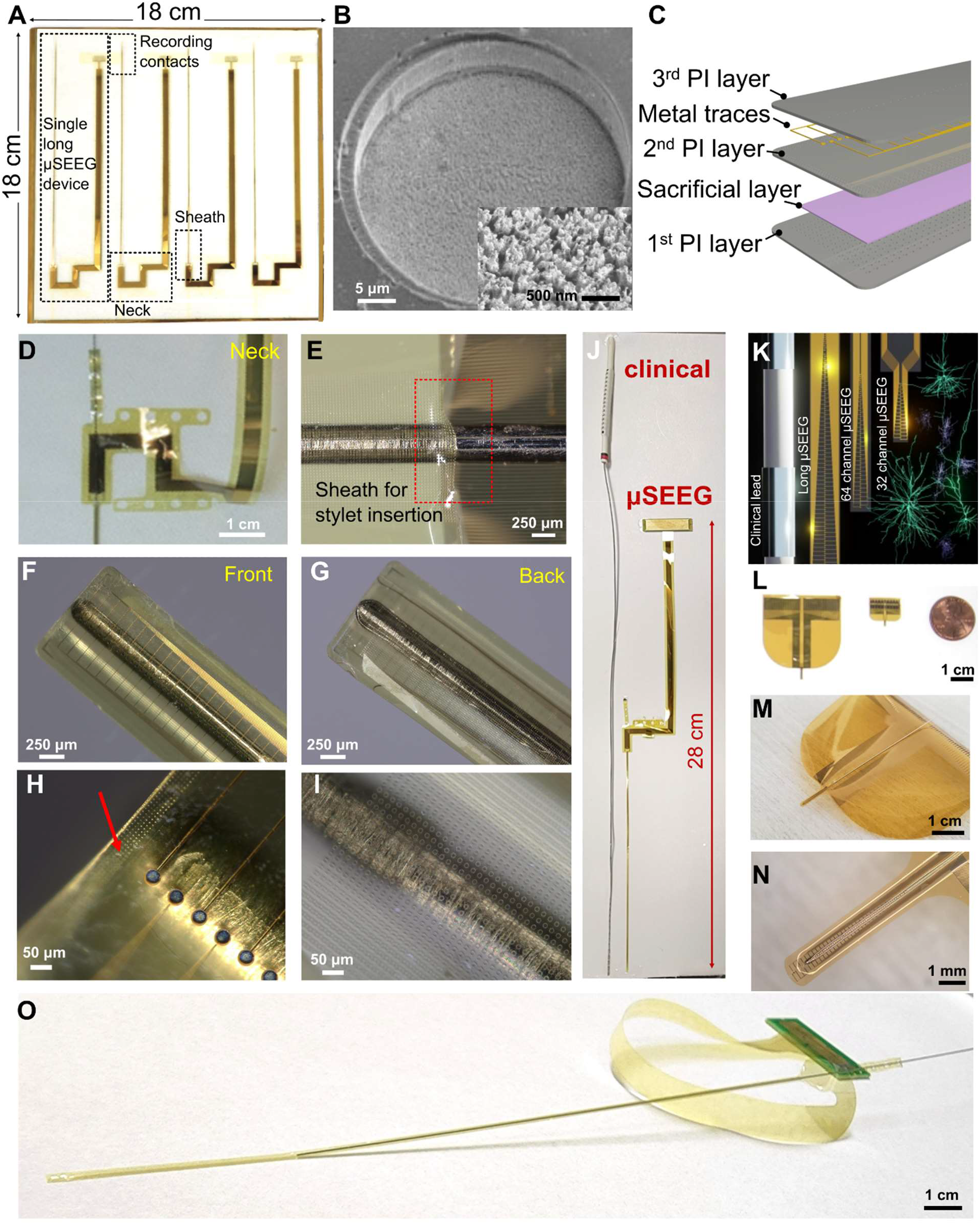
μSEEG electrode arrays. (**A**) Photograph of a single glass substrate plate with four μSEEG electrodes. (**B**) Scanning electron microscope (SEM) image of a single PtNR contact; Inset is a magnified image showing the PtNRs. (**C**) Structural composition of the μSEEG array. (**D**) Photograph showing the ‘neck’ of the array where the U-shape pattern is flipped to provide metal trace extension and circular holes are present to stabilize the inserted stainless-steel stylet. (**E**) Optical microscope (OM) image of the region of insertion of the stylet in the inflatable ‘sheath’ of the μSEEG electrode. OM images of (**F**) front, (**G**) back side of the μSEEG electrode. (**H** and **I**) Magnified OM images at the tip **F** front and **G** back layers. Red arrow indicates holes to enable electrode integrity. (**J**) long 128 channel μSEEG electrode and comparison with a clinical electrode. (**K**) Diagram of the relative scale of human cortical neurons relative to a clinical SEEG lead and μSEEG electrodes (*35-37*). (**L**) Flexibility in manufacturing procedure to produce short 64 channel μSEEG electrodes (left) or short 32 channel μSEEG electrodes (right) and photographs showing (**M**) overall and (**N**) tip of the 64 channel μSEEG electrodes. (**O**) A perspective view of the long μSEEG electrode with partially inserted stylet illustrating the flexibility and slenderness of the electrode body.

## Results

### Manufacturing μSEEG electrodes

To fabricate μSEEG electrodes, we first coated the glass substrate with a sacrificial polyimide layer and followed by the deposition of titanium etch-mask layer that was patterned with circular openings (**fig. S1**). We then coated the glass substrate with two polyimide layers (1^st^ and 2^nd^ PI layer) and an interleaved Ti sacrificial layer (**Fig. 1C**).

When the sacrificial titanium layer is dissolved in a later stage in the process, the two polyimide layers form the structural enclosure (sheath) for the insertion of the stainless-steel stylet. Above the second polyimide layer, we deposited and patterned the metal trace layer with 10 nm/250 nm chromium/gold stack (deposited and patterned twice for a total trace thickness of 520 nm) followed by the deposition of a film of platinum-silver alloy used for the formation of PtNR contacts. This was followed by a top-most polyimide layer (3^rd^ PI layer, **Fig. 1C**) coating. We next induced holes in the 3^rd^ PI layer to expose the platinum-silver alloy films. The shape of the electrode and additional larger holes (**Fig. 1D**) were then etched into the polyimide layer for mechanical stabilization of the stylet. A nitric acid (HNO_3_) etch at 60 ° C dissolved the silver from the platinum silver alloy and exposed the PtNR contacts (**fig. S1**). The resulting structure is then peeled off from the substrate, flipped, and temporarily adhered to another host glass substrate. At this point, the very first sacrificial polyimide layer was etched by O_2_ plasma exposing the titanium etch-mask layer that was pre-patterned with circular openings. Continuation of the O_2_ plasma etching through the circular openings drilled through the 1^st^ PI layer and exposed the titanium sacrificial layer underneath. A final buffered oxide etching dissolved the titanium layers (both sacrificial and etch-mask layer) after which the device is rinsed with flowing de-ionized water.

The stylet is inserted through the mechanical stabilization holes (**Fig. 1D**) and the sheath formed by the two polyimide layers (**Fig. 1E**) to the tip of the electrode (**Fig. 1, F and G**; stylet insertion process illustrated in **fig. S2**). At the very tip the electrode, an array of holes was etched in the 1^st^ PI layer around the sacrificial layer (marked with a red arrow in **Fig. 1H**). As the 2^nd^ PI layer is coated to fill these holes, the interface between the 1^st^ and 2^nd^ PI layers has effectively a larger surface area than a planar one and as a result, better adhesion between the 1^st^ PI layer and the 2^nd^ PI layer is established. The greater mechanical stability afforded by the array of holes prevent the stylet from piercing through the tip when the stylet reaches this interface. The tip of the stylet is mechanically polished to a rounded shape to minimize damage during insertion (**fig. S3**).

The μSEEG electrode was manufactured with a U-shaped neck between the electrode array proper and a continuation of the thin film providing additional length for the metal traces. Once straight edges of the U-shaped electrode are flipped, the total length of the μSEEG electrode becomes 28 cm, on par with the length of a standard clinical sEEG electrode (**Fig. 1J**) but with a total thickness of approximately 15 μm. Overall, the μSEEG electrode after the stylet insertion had ∼1/10 the cross-sectional area of a typical clinical depth electrode while matching its length and its ability to reach to deep brain structures (**Fig. 1J**).

### μSEEG electrodes are robust to tearing and can be implanted and extracted without deformation, producing less damage than clinical electrodes

As these devices must be robust for longer-term implant periods, mechanical strength and resilience against tear were assessed using pull measurements with both the μSEEG electrode and, to compare with a clinical lead, on a 1.2 mm diameter PMT depth electrode anchored on two polyurethane tube regions around a Pt contact. The tensile strengths (critical forces) were 1 MPa (16 mN) for the μSEEG electrode and 14 kPa (48 mN) for the PMT electrode (**fig. S4**).

Since electrode and contact integrity should also be maintained during implantation μSEEG with the stylet without any deformation or loss of function, an acute implantation was first assessed on a phantom brain model. The displacement of a μSEEG and the surrounding phantom brain medium before and after stylet extraction was less than 10 μm (**fig. S5**, N = 6). Electrochemical impedance spectroscopy before and after stylet insertion showed relatively stable 1-kHz impedances, changing from 33.0 ± 2.5 kΩ to 35.0 ± 3.7 kΩ (**fig. S6**), indicating that there was no substantial damage to the device during stylet insertion. The electrodes were extracted in these phantom experiments and all animal and human experiments without any mechanical deformations or tears.

Finally, to test the amount of tissue damage caused by these devices, we implanted rats with one chronic μSEEG electrode with 1.89 mm recording length on one hemisphere and a clinical electrode on the other hemisphere for 14 days (N = 1 electrode). Insertion of the μSEEG electrode resulted in decreased astrocyte scarring, as measured by significantly lower GFAP intensity as compared to the clinical electrode (**fig. S7**). Within 100 μm from the electrode, for example, GFAP intensity in five randomly-placed 100 μm^2^ boxes was significantly lower for the μSEEG electrode (1842 +/-53 a.u.) compared to the clinical electrode (3840 +/-339 a.u.; two-way ANOVA, F (1,24) = 85.93; P<0.0001; Sidak’s multiple comparisons posthoc, p<0.001). In addition, significantly fewer neurons were observed between 0-100 μm from the electrode for the clinical electrode compared to the μSEEG (2.4 +/-1.2 vs. 19 +/-2.3 cells; 2-way ANOVA, F(1,24) = 23.26, P<0.001; Sidak’s multiple comparisons posthoc, p<0.0001). We also imaged the PtNR μSEEG electrodes upon extraction from the NHP brain and observed minimal changes compared to non-implanted ones demonstrating the stability of the μSEEG electrode in tissue (**fig. S8**, N = 3).

### μSEEG flexible design is scalable for multiple acute and chronic applications

To demonstrate the flexibility in the manufacture, design, and use of μSEEG to record neurophysiologically relevant neural activity in multiple settings and species, we tested devices with working neural recording lengths ranging from 1.89 to 7.65 mm, made from either parylene C or polyimide, with microelectrode contacts composed of either PEDOT:PSS or PtNRs (**Fig. 1K; fig. S9; table S1, S2**). We transitioned to all polyimide PtNR μSEEG electrodes after we observed that parylene C PEDOT:PSS μSEEG develop cracks in the parylene C layers and in the PEDOT:PSS layers after stylet insertion whereas polyimide μSEEG did not suffer from any cracks. Additionally, PtNRs contacts did not suffer any delamination from the μSEEG whereas PEDOT:PSS suffered from delamination after stylet insertion in a substantial subset of electrodes, therefore reducing product yield.

All designs used have microelectrode contacts (also called channels, each 30 μm contact diameter for PtNRs and 20 μm contact diameter for PEDOT:PSS) with a center to center spacing of 60 μm (**Fig. 1K, fig. S9; table S2**). We created two short versions: 1) a short 64 channel μSEEG; 2) a short 32 channel μSEEG. The short 64 channel μSEEG includes 64 microelectrode contacts along a recording length of 3.80 mm. Side flaps are incorporated to help with stabilization of the array upon insertion. This design is intended for use in the intraoperative setting and resembles other microelectrode arrays (often called laminar arrays) which were designed to capture activity across the cortical layers (*38*). (**Fig. 1, K to N; figs. S5 and S6; table S2**). The architecture of this system is formatted for use in smaller animals or in recording from the neocortex of humans or larger animal species – such as for use in a brain computer interface. The short 32 channel μSEEG (**Fig. 1, K and L**, right; **fig. S9; table S2**) includes 32 microelectrode contacts along a recording length of 1.92 mm intended for use in chronic recordings in smaller animals.

We also made a longer version designed for accessing deeper structures (simultaneously with lateral cortex) in larger animals. This long μSEEG includes 128 microelectrode contacts at a 60 μm center to center spacing along a recording length of 7.65 mm at the tip of the entire array. This configuration most closely resembles clinical depth electrodes (**Fig. 1, J and O; fig. S9**) although the spacing of the contacts or the incorporation of contacts with diameters larger than 100 μm in a future μSEEG design can be varied for specific end use.

### μSEEG electrodes record local field potential events both acutely and chronically

To test the capabilities of the μSEEG electrode in capturing relevant neural activity (*39*) we recorded from the rat barrel cortex in both the acute and chronic settings. Acute recordings from rat S1 cortex under anesthesia were performed with both a surface μECoG array (*30*) and the 64 channel μSEEG (**Fig. 2A to C; figs. S6 and S10**). When contralateral whiskers were selectively deflected by a directed air puff stream, we found LFP voltage responses (z-scored relative to 0.5 sec before stimulus delivery) and increases in high gamma power (HGP; power between 65 and 200 Hz) on both the μECoG and μSEEG (**Fig. 2D; figs. S11 to S14**). At different depths along the μSEEG electrode and at different channels in the μECoG grid, whisker deflection induced significantly greater LFP and HGP responses than baseline (0.5 sec before stimulation; Wilcoxon rank sum test; p<0.001; n = 39 trials), with some deflections showing reversals in voltages along the depth electrode, also reflected in the current source density (CSD) analysis (**Fig. 2E; fig. S11 to S13**). Further, we found sensory specificity in the responses, with stronger neural responses (in the LFP, HGP and CSD) with stimulation closer to D3, C3, and even E3 (**Fig. 2E; figs. S11 and S12**), allowing us to estimate the location of the electrodes relative to columns of the barrel cortex. The concurrently implanted μECoG surface microelectrode, used to confirm we were recording from the barrel cortex, reflected similar D3, C3, and even E3 whisker-selective voltage and HGP dynamics in response to sensory stimulation (**figs. S13 to S14**).

**Fig. 2.**
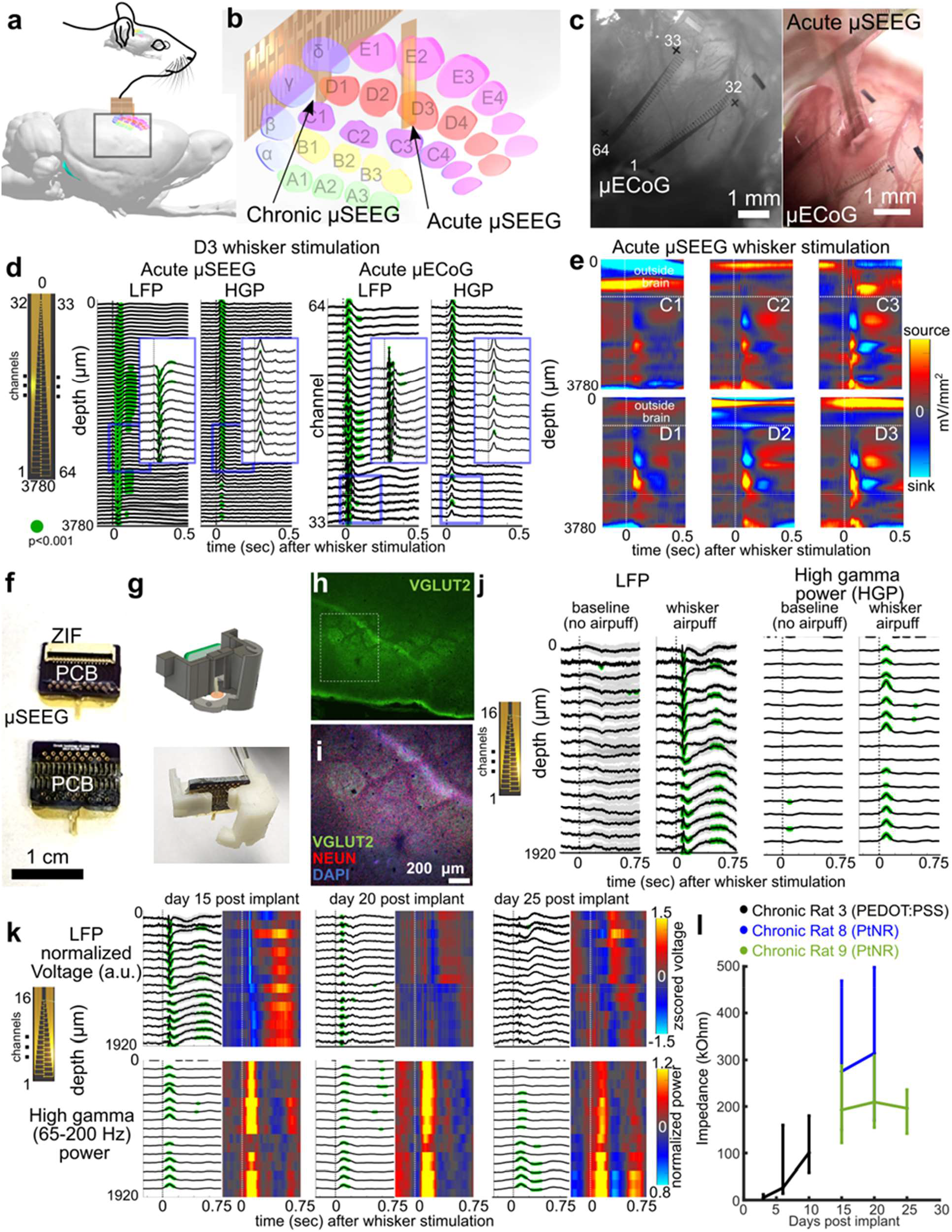
μSEEG electrodes can be used for acute and chronic implantations and recordings. (**A** and **B**) Location and 3D reconstruction of possible locations of the acute and chronic implantation of μSEEG electrode devices for recording from the rat barrel cortex (*40*). (**C**) Images of the implanted μECoG electrode (left) and the μSEEG electrode (right). Note some contacts are outside brain tissue on the μSEEG electrode. (**D**) Example voltage responses across the μSEEG electrode (left) and the μECoG electrode (right) to whisker stimulation at different whisker locations, with the insets zoomed-in views of the voltages and HGP. Green dots indicate significantly different from 0.5 sec before (which is baseline) air puff stimulation to the whisker (Wilcoxon rank-sum test) per channel and across trials. Number of trials >10. (**E**) Increasing responses as air puff stimulation is closer to the C3 and D3 whiskers as indicated by the CSD. (**F**) 32 channel μSEEG electrode for chronic rat recordings and a custom printed circuit board (PCB) with zero-insertion-force (ZIF) connector which electrically connects the device to the recording system via flexible flat cable (FFC). (**G**) 3D-printed headstage for the 32 channel μSEEG electrode. (**H-I**) Electrode location localization as visualized using histology. (**J**) Example voltage and high gamma power (65-200 Hz) responses without stimuli (baseline) versus with stimuli (whisker air puff). (**K**) Responses to air puffs at three different time points post implant in one rat. (**L**) Impedance measures at 1 kHz across multiple days and multiple rats; vertical bars are standard deviation from average values.

After confirming that μSEEG electrodes could detect sensory stimulation-induced neural activity acutely, we developed a 3D-printed headstage for a chronic implantable version of the short 32 channel μSEEG electrode (**Fig. 2, F to G; fig. S15**). We implanted the device successfully in nine rats (**table S1**) with rat barrel cortical responses in three of the nine rats with histological confirmation of the electrode location (**Fig. 2, G to K**). We implanted the devices for 25 days and recorded at three or more time points following implantation to test recording quality and impedance changes over time (**Fig. 2, F to L**). Impedance fluctuated across days per rat but was still low enough to record voltage responses (63.5 ± 49.1 kΩ, 268.0. ± 115.2 kΩ, and 198.0 ± 48.7 kΩ for rat 3, 8, and 9, respectively) across rats and across days post implant (**Fig. 2L**). We recorded voltage responses and changes in high gamma power with whisker stimulation, which was not evident when performing sham controls (trials with no air puffs; **Fig. 2J**). Further, we observed similar voltage responses and HGP recorded by functional microcontacts across the days in individual rats (**Fig. 2K**). This result was confirmed by calculating the correlation between averaged responses across days per channel. In particular, we found that activity during whisker stimulation was more correlated per channel across days (Pearson’s linear correlation average rho across channels: 0.2 ± 0.06 (std), maximum average: 0.73 ± 0.10 (std)) compared to sham controls (Pearson’s linear correlation average rho across days per channel: 0.17 ± 0.03 (std), maximum average: 0.56 ± 0.13 (std)). These differences were also reflected in the high gamma power measures (Pearson’s linear correlation average rho across days per channel: whisker stimulation: 0.33 ± 0.28 (std), maximum average: 0.70 ± 0.15 (std); sham controls: 0.21 ± 0.07 (std), maximum average: 0.54 ± 0.04 (std)).

### μSEEG electrodes acutely record stimulus and anesthesia-induced dynamics across species

Demonstrating that μSEEG electrodes can be used to record clinically relevant dynamics in the human brain requires both scaling up the devices for use in recording from larger brains as well as demonstrating that μSEEG electrodes record clinically and neurologically relevant neural dynamics (*41, 42*). A major goal was to test the μSEEG while modeling settings and paradigms which could be used in acute or chronic clinical mapping of activity (*6, 43*). Therefore, we recorded neural activity using the short 64 channel μSEEG in three different settings: 1) from the somatosensory cortex in an anesthetized pig, 2) in an anesthetized NHP in the operating room, and 3) in the operating room from human cortex preceding tumor resection (**Fig. 3, fig. S9; table S1**).

**Fig. 3.**
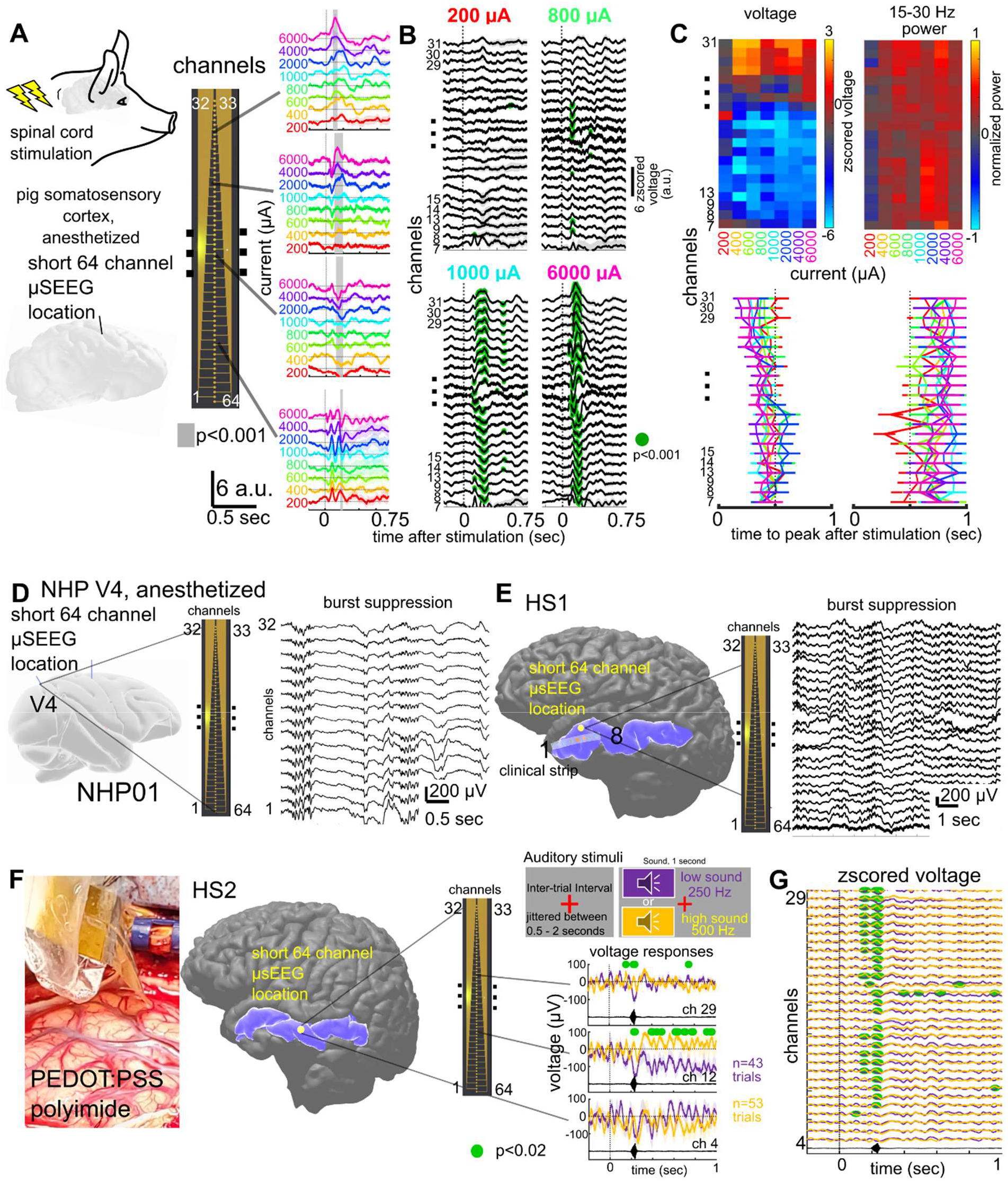
Stimulated neural activity and ongoing clinically relevant neural dynamics can be recorded acutely using the short 64 channel μSEEG electrode. (**A**) Direct electrical stimulation of the spinal cord while doing an acute short μSEEG electrode recording from the pig cortex, with responses increasing with increasing injected current. Grey bar indicates significantly different between current steps, Wilcoxon rank sum test. (**B**) Voltage responses along the electrode depth with more responses significantly different to baseline (0.5 sec before stimulation) occurring more with higher current levels (green dots, Wilcoxon rank sum test). (**C**) Top: Two-dimensional heat map of the largest voltage deflection from baseline per channel and per current step (left) and the maximum peak in beta (15-30 Hz) power oscillations (right). Bottom: time to peak voltage (left) and peak beta band power (right) after stimulation per channel and current step. Grey dotted line indicating 0.5 seconds after stimulation. (**D**) Acute implant of the short μSEEG electrode into V4 in an anesthetized NHP and ongoing evidence of burst suppression. (**E-G**) Acute implantation of short μSEEG electrodes into left anterior temporal lobe middle temporal gyrus (highlighted in blue) to be resected in the course of treatment of epilepsy in two participants, HS1 and HS2 with a photograph of the implant, a three-dimensional reconstruction of each participants’ brain and the relative location of the μSEEG electrode (yellow dot) with a zoomed in inset view of the 64 channels as implanted. The polymer substrate was parylene C for HS1 and polyimide for HS2 with the contacts made of PEDOT:PSS. (**E**) Spontaneous ongoing activity with burst suppression along the electrode depth. (**F**) Auditory responses to low and high tones presented at random in sequence with varying jitter times while recording activity in the lateral temporal lobe (pictured, see **Materials and methods**). Left: pictured inserted electrode and placement in the brain. Top right: Stimulus presented. Bottom right: Voltage responses at different depths averaged across trials for three channels. Green dots indicating p<0.02 significant difference between low and high tones, Wilcoxon rank sum test. (**G**) Differences in the responses which varied across the depth of the electrode. Z-scored voltage responses at multiple channels at different depths averaged across trials. Green dots indicating p<0.02 significant difference between low and high tones, Wilcoxon rank sum test.

Intraoperative clinical mapping often involves the use of stimulation to delineate functional (eloquent) tissue and connectivity. To model this paradigm, we stimulated the pig spinal cord with a bipolar stimulator and recorded with the μSEEG in the pig cortex to map responsiveness and connectivity. We recorded changes in neural activity across cortical layers in the somatosensory cortex induced by direct electrical stimulation in the spine using the short 64 channel μSEEG electrode, stimulating with currents ranging from 200 μA to 6000 μA (**Fig. 3, A to C**). We found significantly increasing voltage responses with increasing injected current per channel (p<0.001; Kruskal Wallis test; **Fig. 3A, fig. S16**). The response waveforms varied between the different contacts, including a field reversal approximately in the middle of the implanted electrode. This field reversal was most obvious at stimulation currents > 800 μA (green dots, **Fig. 3B**, significantly above a baseline taken 0.5 sec before stimulation, Wilcoxon rank sum test). When we plotted the largest absolute voltage deflections from baseline, we found a clear division in the voltage between the deeper contacts and the superficial contacts. This high-resolution laminar distribution of the responses also was reflected in differences in oscillatory power across the cortical layers. We found gamma (30-55 Hz; p= 0.0056 for 6000 μA) power in the more superficial contacts (Kruskal-Wallis multiple comparisons test; **Fig. 3C, fig. S16**). We also found response timing differences, with the time to voltage peak and HGP peak shorter in middle and superficial contacts (resulting in a Pearson’s linear correlation between peak timing and channel number: voltage-rho= -0.11; p<0.0001; HGP-rho= -0.04; p=0.03) with the trend reversed with the peak beta power (beta power-rho= 0.04; p=0.03; **Fig. 3C, fig. S16**). In other words, the μSEEG electrode recorded stimulation-induced activity with cortical layer-specificity at a spatial resolution (60 μm contact to contact pitch) not possible with current clinical leads (resolution on the scale of millimeters; **fig. S9**).

In a second intraoperative paradigm, we recorded neural activity across cortical layers in the visual cortex of an anesthetized NHP using the short 64 channel μSEEG electrode. We found ongoing anesthesia-related burst suppression which could be detected using automatic approaches along the depth electrode ((*44*); **Fig. 3D, fig. S16**). As shown in previous preparations and other species, we found a gradient of activity across the array, with more detected bursts early in the recording and even relative to suppression epochs (as represented by the burst-suppression ratio) in more superficial contacts ((*44*); **fig. S16**). This resulted in high negative correlation values between electrode depth into the tissue and detected bursts over 300 seconds (r= -0.73; p = 0.0021; Pearson’s linear correlation; **fig. S16**).

Finally, in a true test of the translational feasibility of the μSEEG, we acutely implanted short 64 channel μSEEG electrodes in the left middle temporal gyrus in two separate human patient participants (**Fig. 1F; Fig. 3, E to G**) undergoing temporal lobe resection for clinical reasons. With each participant, we inserted a single 64 channel short μSEEG devices into tissue which the clinical team determined would be resected. The recordings were brief (10 minutes) yet we were able to record ongoing spontaneous activity. In one case the participant was under general anesthesia and there was clear evidence of anesthesia-induced burst suppression in the recordings (HS1), also detected through an automatic algorithm ((*44*); **Fig. 3E**). Like in the NHP, the number of detected bursts was increased in more superficial contacts, resulting in a correlation between depth (into the tissue) and burst detections of r= -0.5927 (p = 0.0023; Pearson’s linear correlation, over 300 seconds of recording).

In the second recording, the participant was awake under monitored anesthesia care (MAC) and listened to low and high auditory cues (HS2; see **Materials and Methods**; **Fig. 3, F and G**). The neural responses were significantly different between low and high tones in the z-scored voltage values and in HGP at the onset of the sound (p<0.02, corrected for multiple comparisons; Wilcoxon rank sum test). Further, there were more significant differences in the responses between low and high sounds in superficial array contacts (**Fig. 3G**).

### μSEEG electrodes detect single unit cortical activity

A key purpose of the μSEEG electrode is to offer advantages over current clinical depth electrodes including increased spatial resolution as well as increased neural resolution. To test whether the μSEEG device can record neural activity at multiple depths in the brain closer to the scale of the human brain, we designed and built the long μSEEG electrode (**Fig. 1 and 4**). The 128 channel long μSEEG electrode was built to most resemble clinical depth electrodes with a working recording length of 7.65 mm at the tip of the electrode that is, overall, 28 cm long, 1.2 mm wide, and 15 μm thick which would allow insertion and recording from deeper brain structures. The electrode contacts in this design are concentrated at the tip of the device with inter-contact distances of 60μm. To test if we could record single neuron activity at depth, we recorded neural dynamics in an NHP which was awake but resting and viewing flickering light emitting diodes (LEDs) (*29*) to test for visual responses. The long μSEEG electrode was held by a microelectrode microdrive (see **Materials and Methods; fig. S17**) to drive the microelectrode to multiple depths from the surface of the cortex within an implanted chamber (**Fig. 4A**). Along the trajectory moving toward the thalamus, we stopped and recorded at three different depths to examine spiking activity in the cortex as well as in white matter (**Fig. 4B**). At depths 1 and 2, we found we could record spikes which clustered into single unit and multi-unit activity (MUA) (depth 1: 1 MUA cluster, 4 single unit clusters; depth 2: 5 MUA clusters, 31 single unit clusters; **Fig. 4, C to E**), which we determined by examining the autocorrelation of the spike times and the waveform consistently through time using Kilosort (*45*). We did not find any identifiable single unit activity at depth 3, we were likely mostly in white matter at that depth (depth 3: 13 MUA clusters; **Fig. 4, F to G**). The units and MUA clusters were distributed at different distances and locations along the 128 contacts of the long μSEEG with a range of spike rates, most of which were around 2 Hz (**Fig. 4H**). Finally, we found the waveform measures show that the units sampled at depths 1 and 2 were clustered in amplitude, the peak-trough ratio, and spike duration measures compared to the MUA clusters found at depth 3 (**Fig. 4I**). In other words, these clusters are more likely single unit activity or putative neurons since they were detected while the recording contacts were in cortex but not while in white matter.

**Fig. 4.**
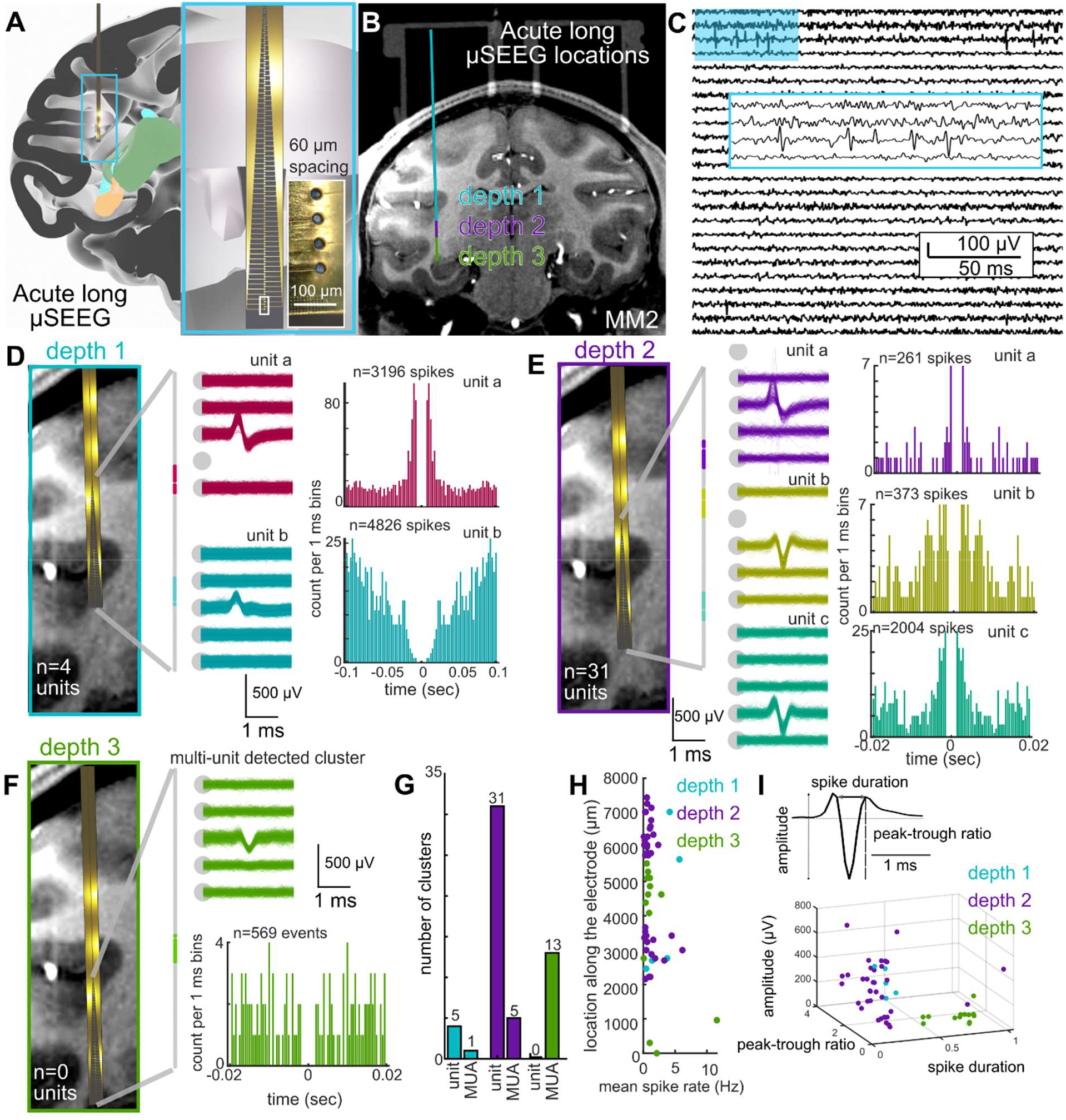
Single unit activity could be recorded using long μSEEG electrodes. (**A**) Three-dimensional reconstruction of the locations of the long μSEEG inserted at multiple depths into the parietal lobe, temporal lobe, and deeper into the tissue, with a zoomed-in view of the microelectrodes in the long μSEEG (*46, 47*). (**B**) MRI with the overlaid CT (chambers above) and the three putative long μSEEG depths in the brain. (**C**) Example recording from the second depth to show single unit spiking activity (filtered to between 300 and 6000 Hz). (**D** and **E**) Example Single units recorded at electrode depth 1 (D) and depth 2 (E) showing overlaid waveforms and the autocorrelation of the spike times. (**F**) Single unit activity was not observed at depth 3 which seemed to be in white matter, but possible MUA was recorded at depth 3. (**G**) Numbers of detected single units and MUA clusters. (**H**) Location detected clusters relative to the mean spike rates for the different depths. Each dot is a cluster (which can represent single units or MUA). (**I**) Spike waveform measures of single units and MUA waveforms showing separation of events detected at depths 1 and 2 versus 3.

## Discussion

We developed a μSEEG electrode that is implanted with a stylet inserter similar to clinical SEEG electrodes, but can be tailored in its range of depth to sample cortical and or deep structures in the brain (or both). This new μSEEG electrode can record broadband spontaneous and evoked neurophysiological activity including LFP, CSD and single/multi-unit activity across a variety of species including humans and across entire depths of the brain. While the μSEEG construction is robust, it induced less apparent tissue damage than clinical SEEG electrodes. The layout, shape, and size of the μSEEG electrode could be generated with customizable designs (**Fig. 1, K to N**) by leveraging established display screen fabrication techniques on large glass wafers. Fabrication on glass wafers also promises excellent scalability. Glass panels used in the manufacture of displays use plates a few square meters in area indicating that large number of arrays can be manufactured even if the arrays are long. Furthermore, the high-resolution of lithographic capability can achieve 1.2 μm for both metal line and space (L/S) of flat panel displays (*48*). Therefore, the number of contacts can be increased well beyond 128 channels presented here importantly afforded at a much lower manufacturing cost that clinical and other research depth electrodes. A 240 sq. in. monitor has a retail price of nearly a $100 with active transistors and light emitting diodes. The same area can be used to manufacture at least 20 μSEEG electrodes, pointing to significantly lower costs than current clinical SEEG electrodes (> $1,000 per electrode) when manufactured at scale. This cheaper new manufacturing approach and the added spatial resolution and sensitivity to cellular activity in a smaller form factor can advance our ability to study and treat the human brain and will help broaden the access of the technology to underserved communities and other brain diseases.

One potential limitation of these designs is cross-talk amongst the channels. While we have not definitively quantified cross-talk in the recordings, we observed a strong common-mode signal on all contacts that we subtracted in order to delineate CSD dynamics. If inter-channel cross-talk is a substantial issue, future designs could involve distributing the metal traces in separate polymer layers. Another limitation concerns connectorization. Current connectors do not match typical clinical standards. Improving the back-end of the devices is an area of active development. Further, the current design includes contacts facing only one direction along the electrode length. Future designs can involve developing multiply directional contact sampling.

Nevertheless, advantages include the size and shape of the electrodes as well as the capabilities of the devices. For instance, the width of the μSEEG was still destructive to brain tissue, unlike ultraflexible nanoelectronic probes (*49*). However, a human-grade electrode that can reach 10 cm deep into the brain with 128 contacts necessitated the stylet-guided μSEEG design, especially with stimulation considerations where the metal traces need to be sufficiently wide to reduce serial resistances and associated potential drops that compromise the long-term electrode stability. Lastly, the μSEEG can also offer stimulation with favorable stimulation characteristics with PtNRs compared to clinical SEEG electrodes. We prepared PtNR electrode contacts with 1 mm diameter to test how PtNR compares with clinical electrodes in delivering stimulation in saline (**fig. S18**). In addition to manufacturing variable contact sizes with this approach, the PtNR contacts at a 1 mm diameter offer smaller voltage transients and higher electrochemical safety limits when delivering direct electrical stimulation than clinical electrodes of similar surface areas (**fig. S18**). Thus, the μSEEG can offer micro and macro stimulation capability for use in future deep brain stimulation (DBS) or direct electrical stimulation application.

In conclusion, the new μSEEG electrodes provide an innovative approach enabling recording across the entire depth of the human brain with greater resolution than ever achieved before. The smaller volume of the μSEEG electrode and its compatibility with procedures used in clinical practice paves the way to increasing our understanding of brain diseases and offering novel and clinical interventions.

## Acknowledgments

We are grateful for the technical support from the nano3 cleanroom facilities at UCSD’s Qualcomm Institute where the depth electrode fabrication was conducted. This work was performed, in part, at the San Diego Nanotechnology Infrastructure (SDNI) of UCSD, a member of the National Nanotechnology Coordinated Infrastructure, which is supported by the NSF (grant ECCS1542148). We thank Yangling Chou, Aaron Tripp, Fausto Minidio, Daniel J. Soper, and Alexandra O’Donnell for their help in collecting the data.

## Funding

National Institutes of Health BRAIN® Initiative UG3NS123723-01 (S.A.D.)

National Institutes of Health BRAIN® Initiative R01NS123655-01 (S.A.D.)

National Institutes of Health NBIB DP2-EB029757 (S.A.D.)

National Institutes of Health F32 postdoctoral fellowship MH120886-01 (D.R.C)

National Science Foundation Award no. 1728497 (S.A.D.)

National Science Foundation CAREER no. 1351980 (S.A.D.)

National Science Foundation Graduate Research Fellowship Program no. DGE-1650112 (A.M.B.)

MGH - ECOR (S.S.C.)

K24-NS088568, R01-NS062092 (S.S.C.)

Tiny Blue Dot Foundation (to S.S.C. and A.C.P.)

NIH NS047101 to the UCSD Neuroscience Microscopy Core

National Institutes of Health BRAIN® Initiative K99 NS119291 (K.J.T.)

## Author contributions

Conceptualization: SAD, EH, SSC, KL, ACP, YGR

Methodology: KL, ACP, YGR, DRC, KT, YK, JP, YT, JL, AMB, RV, JRM, SMR, JCY, AB, RMR, ZMW, SIF, AMR, SBH, EH, SSC, SAD

Investigation: KL, ACP, YGR, DRC, KT, YK, JP, YT, JL, AMB, RV, SMR, JCY, AB, RMR, ZMW, SIF, HSU, EH, SSC, SAD

Visualization: KL, ACP, YGR

Funding acquisition: SAD, EH, SSC

Project administration: SAD, SSC

Supervision: SAD, EH, SSC, JP, MR, ZMW, SIF

Writing – original draft: ACP, KL, SAD, YGR

Writing – review & editing: KL, ACP, YGR, DRC, KT, YK, JP, YT, JL, AMB, RV, JRM, SMR, JCY, AB, RMR, ZMW, SIF, HSU, AMR, SBH, EH, SSC, SAD

## Competing interests

The authors declare the following competing interests: KL, YGR, and SAD and the University of California San Diego filed a patent application for the manufacture of the novel depth electrodes. YT, AMR, and SAD have competing interests not related to this work including equity in Precision Neurotek Inc. and SAD in FeelTheTouch LLC. SAD was a paid consultant to MaXentric Technologies. AMR has an equity and is a cofounder of CerebroAI. AMR received consulting fees from Abbott Inc and Biotronik Inc. The MGH Translational Research Center has clinical research support agreements with Neuralink, Paradromics, and Synchron, for which SSC provide consultative input. The other authors declare that they have no competing interests.

*Data and materials availability*: All data obtained in this study are either presented in the paper and the Supplementary Materials or deposited in open database. Animal brain recording data could be accessed at OpenNeuro (https://openneuro.org/), and the human brain recording data could be found in Data Archive BRAIN Initiative (DABI) (https://dabi.loni.usc.edu/) using the iEEG BIDS format (*50*). Custom MATLAB code (version R2021a) are available in GitHub.

## Supplementary Materials

Materials and Methods

Supplementary Text

Figs. S1 to S18

Tables S1, S2

References (*51*–*66*)

## Notes

### Summary of Updates

Figure 2 had an update.

