## Supplemental Methods and Figures for "Flexible, Scalable, High Channel Count Stereo-Electrode for Recording in the Human Brain" (from another version of this preprint: version 1, 2022-11-09)

**This PDF file includes:**

Materials and Methods
Figs. S1 to S18
Tables S1 to S2

### Materials and Methods

#### 1. Study design

Objectives of the study were to: (i) demonstrate implantable thin-film electrodes (µSEEG) produced using reproducible, customizable, and high throughput manufacturing pipeline to be implanted in the brain such as in the operating room using similar brain implant techniques to standard clinical depth electrodes, (ii) use the electrode to record neurophysiologically relevant neural activity in multiple settings and species and; (iii) reach deep brain structures with high spatial sampling resolution and channel count with a much thinner electrode body. Following mechanical and electrochemical characterization of the properties of the new µSEEG devices, we recorded neurophysiological activity in four species (rat, pig, rhesus macaque, and human) under general anesthesia, while awake or under monitored anesthesia care (MAC; **table S1**). The primary goal in the physiological recordings was to use sensory (auditory or mechanosensory) or electrical stimulation as well as spontaneous neural activity to determine if the µSEEG devices can record known neurophysiological signatures, whether in the local field potential or the single unit spiking activity (action potentials). All rodent and pig experiments were approved by the UCSD Institutional Animal Care and Use Committee (IACUC) (**table S1**). Experimental procedures on rhesus macaques (N=2) were carried out in accordance with the Guide to the Care and Use of Laboratory Animals and approved by the IACUC at Massachusetts General Hospital (MGH; **table S1**). We successfully gathered depth recordings of all intended stimulus-evoked activity which included baseline recordings and at least 10 trials per condition (whether injected current level or sensory stimulation). Endpoints for stopping recordings and experiments were determined by the IACUC protocols. For the NHP recordings, we planned to limit µSEEG to 15 minutes-30 minutes for lowering and recording, particularly under anesthesia, to minimize the duration of the case.

Two human participants were recruited to participate in this study at MGH under the Partners Institutional Review Board (now the Mass General Brigham Institutional Review Board which covers MGH). The possibility of conducting research recordings were only discussed with each patient after the decision to proceed with the surgery had been made and following consultation with the treating neurosurgeon and clinical team. Further, we did not enroll patients with clearly impaired decision-making abilities. Both participants were involved voluntarily, provided informed consent, and were informed that participation in the experiment would not alter their clinical treatment in any way and that they could withdraw at any time without altering their clinical care. Finally, case selection and the decision to use the penetrating µSEEG could be inserted into tissue which was identified by the clinical team to be resected during the course of normal clinical care. Data from each participant was de-identified prior to analysis using automatic approaches. In both cases, ongoing spontaneous activity was the control acquiring a baseline recording prior to or throughout the given recording period. Trial numbers for the second participant for the auditory stimulation task within a given subject recording were determined based on a maximum recording time of 15 minutes in the OR setting. We did not include a randomization of subject selection in this study because each recording from each individual was considered a separate dataset. Experimental conditions and a summary of all subjects are presented in **Table S1**.

Inclusion and analysis of data per experiment was decided based on the following criteria: 1) histological or physiologic confirmation that the electrode was in the targeted barrel cortex in the rodent studies; 2) if the noise level in the recording was low enough to examine neurophysiological signatures. In the former case, we had to reject 6 of 9 rodent implants as the implanted electrodes were not in the rodent barrel cortex. In the latter case of neural activity and noise levels, we were forced to exclude analyses of 2 of the 3 pig recordings due to overwhelming electrical noise. Both the tests performed with humans and non-human primates (NHPs) were included, therefore passing these criteria. Following preprocessing, trial rejection included removing trials with wayward voltage deflections >6 standard deviations from the baseline activity. Overall, only a maximum of 2 or 3 trials were removed per condition across the data sets. These data were not blinded, and we did not include a randomization of subject selection in this study. We did not use power analyses as this particular study involved observational approaches to test the functionality of the devices

### 2. Fabrication of electrodes

#### 2.1. Long µSEEG fabrication

**Figure. S1** summarizes the fabrication process of the µSEEG electrode. Polished and cleaned photomask-grade soda lime glass plates (Nanofilm) with dimensions of 7” × 7” × 0.06” were used as substrates for the fabrication. First, a Micro-90 (International Products Corporation) diluted with deionized (DI) water (0.1%) layer was spin-coated on the glass substrate as a release layer for the polyimide electrodes in the last step of the fabrication processes. Subsequently, a sacrificial 5-µm-thick-polyimide (PI-2611 from HD Microsystems) was initially deposited by conventional spin-coating, soft-baking, and curing (340 °C for 3 hrs, 3 °C/min in $N_2$ ambient) processes. Then, a 60-nm-thick Ti hard mask for net layer formation was formed on the 1st polyimide layer. The Ti hard mask was proceeded by a standard lithography, descum, metal deposition, and lift-off process using AZ5214E-IR photoresist (MicroChemicals), maskless photolithography system (Heidelberg MLA 150), UV flood exposure system (DYMAX), plasma etcher (Oxford Plasmalab 80), and e-beam evaporator (Temescal). The Ti hard mask contains via patterns with hole arrays with a diameter and a spacing of 5 µm for both, and a 1.5 mm wide and 0.6 mm long rectangular shape for a sheath.

A 1st polyimide layer was spun-coated, soft-baked, and cured at 330 °C for 3 hrs in $N_2$ ambient following pre-curing of the substrate to dry out any solvent or moisture trapped in the polyimide surface under 340 °C for 3 hrs in $N_2$ ambient. The 1st polyimide layer will role as the sacrificial bottom layer with net of holes in the final product. Then, a 60-nm-thick Ti sacrificial layer was deposited using same method for the Ti hard mask preparation. The Ti sacrificial layer also acts as etch-stop layer during the final net formation etching step using $O_2$ plasma. To increase adhesion between the 1st and a 2nd polyimide layer, hole arrays with a diameter and a spacing of 5 µm for both are patterned along the Ti sacrificial layer using AZ 1518 (MicroChemicals). Then, the 1st polyimide layer was selectively etched by $O_2$ plasma (50 mTorr, 200 W for 7 min), and the photoresist layer was removed by solvent cleaning processes.

After processes on the 1st polyimide layer, the glass substrate was baked at 330 °C for 2 hrs in $N_2$ ambient. Then, a 2nd polyimide layer was coated and cured under 320 °C for 2 hrs in $N_2$ ambient. Subsequently, metal traces with a width and a spacing of 3 µm for both were formed on the 2nd polyimide layer. Metal traces were composed of Cr/Au (10/250 nm) and the entire lithography, deposition, and lift-off process was repeated on top of the first metal lead layer to form Cr/Au/Cr/Au (10/250/10/250 nm) traces. This double-patterning process was employed to increase yield and reduce risk of photoresist particles from compromising the thin traces.

After the double-layer metal leads formation, a 30 µm-diameter-PtAg alloy was formed selectively on the individual micro-contact recording sites by photolithography, descum, and PtAg alloy co-sputtering using the maskless photolithography system with NR9-6000 (Futurrex) photoresist, plasma etcher, and DC/RF magnetron sputter (Denton Discovery 18), respectively. The detailed fabrication methods and characteristics of PtNRs can be found elsewhere (*29*, *30*). Notably, this process involves a selective etching of silver in a dealloying process, leaving behind non-toxic platinum. A 60-nm-thick Ti capping layer was deposited on top of PtAg alloys to prevent oxidation in air or under oxygen plasma in the following processes.

A 3rd polyimide layer was then coated and cured under 300 °C for 1 hr in $N_2$ ambient. On top of the polyimide layer, a 60-nm-thick-Ti hard mask was deposited and AZ5214E-IR photoresist was coated on top of the Ti layer and patterned to define the outline of the electrodes. Then, $SF_6$/Ar plasma was used to selectively etch the Ti layer through the photoresist. To accomplish a 1st deep etch step, 3 hrs of $O_2$ plasma was performed to etch 3rd, 2nd and 1st polyimide layers until exposing the Ti hard mask layer on the sacrificial polyimide layer.

Following the 1st deep etch process, a-8-µm-thick AZ12XT-20PL-10 (MicroChemicals) was spun-coated and patterned to open via holes for recording sites and printed circuit board (PCB) contact pads, and the outline of the electrodes. Then, $SF_6$/Ar plasma was used to etch Ti hard mask layer and expose the surface of the 3rd polyimide layers on the recording sites and the sacrificial polyimide layers along the outline of the electrodes. Subsequently, the exposed polyimide layers were etched by $O_2$ plasma for 1 hr.

To protect polyimide layers from dealloying process to form PtNRs using nitric acid, a 100-nm-Ti was conformally deposited using sputtering system and a 1-µm-thick parylene C was additionally coated using a parylene deposition system (Specialty Coating Systems 2010 Labcoter). To open via holes for recording sites, a-8-µm-thick AZ12XT-20PL-10 was spin-coated and patterned again. Then, $O_2$ plasma and subsequent $SF_6$/Ar plasma was used to etch the parylene C and Ti layers including the Ti passivation and cap layers to expose the PtAg alloys. The glass substrate was immersed into 60 °C nitric acid for 2 min for dealloying of PtAg alloys and rinsed in de-ionized (DI) water. The microscopic morphology of the PtNR is shown in the scanning electron microscope (SEM) images (**fig. 1B**).

The parylene C and Ti passivation layers were removed by $O_2$ plasma and 6:1 buffered oxide etchant (BOE), respectively. Then, the glass substrate was immersed in DI water to delaminate the electrodes from the glass substrate with the dissolution of the underlying Micro-90 layer.
The delaminated electrode was then flipped and transferred onto a carrier glass wafer. Following the flip transfer, the net of hole arrays was formed on the 1st polyimide layer by $O_2$ plasma removing the sacrificial polyimide layer and selectively etch the 1st polyimide layer through the via openings in the Ti hard mask. The etching time was set longer to obtain clean opening of hole arrays. Note that the 2nd polyimide layer was protected, during the over etching procedure, by the Ti sacrificial layer prepared between 1st and 2nd polyimide layers. Lastly, the Ti hard mask and the sacrificial layers were all dissolved by BOE and the electrode was immersed in DI water for ~6 hrs to rinse any residual BOE.

### 2.2. Stylet insertion

Medical grade stainless-steel 316 stylets (Ø 125 µm) were used as a shuttle to guide the µSEEG electrode to a target region. The stainless-steel was chosen as a shuttle material over a tungsten rod or a silicon shuttle because of its hydrophilic surface, which helps to avoid adhesion between the hydrophobic polyimide surfaces (*51*). These stainless-steel shuttles are the standard stylets used for the implantation of clinical depth electrodes. The stainless-steel stylet was mechanically polished to have smooth surface at its tip to prevent mechanical damages of the µSEEG electrode during the stylet insertion (**fig. S3**).

The stylet was inserted through the sheath on the stylet bottom layer and travelled along the space between the stylet upper and bottom layers where the Ti sacrificial layer was etched away. The retractable stylet provides rigidity to the polyimide electrode tip and supports implantation to the cortex without electrode deformation. The stylet insertion procedure is as follows: First, the µSEEG electrode and the stylet was placed on a clean glass substrate (**fig. S2A**). Then, the stylet was inserted through the stitch holes to restrict lateral movement of the stylet and the sheath of the electrode (**fig. S2B**). Subsequently, DI water was sprayed along the electrode strips to open the space between the stylet upper and lower layers. In a few seconds after DI water spraying, the stylet could be travelled along the space (**fig. S2C**) and finally reached to the tip of the electrodes (**fig. S2D**). **Figure S2E** shows scale of the µSEEG electrode after the stylet insertion and comparison with the clinical electrode indicating that a length of the stylet inserted part of the µSEEG electrode is long enough to access to the deep brain structure. A cross-section view SEM image (**fig. S2F**) shows the stylet upper and bottom layers with the stylet in between after recordings from non-human primate (NHP). The tip of the µSEEG electrode was deliberately opened by a razor blade to observe cross-sectional details of the µSEEG electrode with bio-materials remained on.

2.3. Short µSEEG fabrication

The short 64 channel µSEEG electrode has a large space (flaps) near the interconnection metal leads for surgeon's convenience for electrode handling. It is also designed to bond with anisotropic conductive films (ACF) with commercial off-the-shelf ribbon cables to connect the electrode to the external characterization circuitry (**fig. S6A**). The electrode consists of three polyimide (or parylene C) layers to not only passivate metal leads, but also carry the stylet during implantation (**fig. S6B**). Au was used to serve sacrificial layer to separate 1$^{st}$ and 2$^{nd}$ polyimide layers for the stylet insertion by dissolving of Au layer using Au etchant (TFA, Transene Company, Inc.). A metal lead layer of Cr/Pt (20/100 nm) was embedded between 2$^{nd}$ and 3$^{rd}$ polyimide layers, but recording electrodes were via-opened. To enhance recording qualities, PEDOT:PSS was electro-deposited on the Cr/Pt recording electrodes. PEDOT:PSS was electrodeposited from 0.01 M 3,4-Ethylenedioxythiophene (EDOT) in 2.0 g per 100 mL Poly(sodium 4-styrenesulfonate) aqueous dispersion under galvanostatic conditions at a potential of 0.9 V versus Ag/AgCl in a three electrode setup, i.e., Ag/AgCl electrode as a reference electrode, a large Pt electrode as a counter electrode, and the Pt contacts on the probes as the working electrodes, at a constant temperature of 27 °C using a Gamry potentiostat (Gamry Interface 1000E; Gamry Instruments). Polymerization was driven for 20 s at current density of 5 mA/cm$^2$.

After the stylet was inserted in the µSEEG electrode, the large space and bonding pad regions were bent due to strain of the polyimide layers because of its thickness and geometry (**fig. S6C**), but it is confirmed that a region for the recording electrodes remained straight with the stylet (**fig. S6D**) and PEDOT:PSS electrodes, Cr/Pt metal leads, and polyimide layers are not damaged (**fig. S6E** and **F**) . **Figure S6E** also displays a detail of the electrode layout showing hole arrays patterned along the metal leads to help wet etching of the Au sacrificial layer. The diameter and center to center spacing of electrode sites is shown as 20 and 60 µm, respectively (**fig. S6F**), and the electrode consisted of 64 channels along 3.80 mm in a laminar manner. The dimension and the number of electrodes can be adjusted based on the target of merit.

Electrochemical impedance spectroscopy (EIS) of the electro-deposited PEDOT:PSS was performed, in 1X phosphate buffer saline (PBS) solution (**fig. S6G**). Three electrode configuration i.e., PEDOT:PSS electrodes as the working electrode, Ag/AgCl electrode as a reference electrode, a large platinum electrode as a counter electrode was used. 10 mV root mean square (RMS) sinusoidal signal with zero DC bias were applied and the frequency was swept from 1 Hz to 10 kHz using a Gamry potentiostat (Gamry Interface 1000E; Gamry Instruments). Electrochemical impedance at 1 kHz is commonly used as the benchmark for the characterization of neural electrodes, as this frequency corresponds to spiking activity (*52*). The average impedance magnitude of PEDOT:PSS electrodes across 64 channels was 33.0 ± 2.47 kilohms at 1 kHz and their phase indicates their faradaic electrochemical interfaces. After stylet insertion, the average impedance magnitude maintained similar values of 35.0 ± 3.67 kilohms on 64 channels, still lower and consistent contrast to Pt electrodes (**fig. S6H**), indicating that the stylet was successfully inserted between the polyimide layers without any damage to the channel or PEDOT:PSS separation from Pt electrodes.

#### 2.4. Surface µECoG fabrication

Parylene C was chosen as the surface µECoG electrode material because of its superior conformability and hydrophobic surface which makes stable electrical and mechanical contact with the surface of the cortex (*32*). The µECoG electrode is designed to perform dual column recordings while the µSEEG electrode measures intracortical signals between the columns (**fig. S10A and B**). It is also designed to bond with ACF with commercial off-the-shelf ribbon cables to connect the device to the external characterization circuitry (**fig. S10A**). The fabrication process of parylene C µECoG electrode was reported elsewhere (*29, 30, 33*).

First, an anti-adhesion layer, Microo-90 diluted with DI water (0.1 %) was spun-coated on a 4” Si carrier wafer to enable parylene C device release from the Si carrier wafer. A 1$^{st}$ parylene C layer (3 µm) was deposited on top of the anti-adhesion layer by chemical vapor deposition (Specialty Coating Systems 2010 Labcoter). A Cr/Au/Ti (10/100/50 nm) metal lead layer was evaporated and patterned via lift-off process. Cr layer serves as an adhesion layer between Au conduction layer and parylene C. Ti layer was deposited to protect Au layer during later via dry-etching process. A 2$^{nd}$ parylene C insulation layer (3 µm) was deposited. A Ti (50 nm) layer was deposited and patterned via lift-off process on the second parylene C layer as an etch hard mask against O2 plasma etching. The Ti hard mask was patterned to define 1) via openings of the Au electrode sites and bonding pads, 2) shape of the device body and 3) perfusion holes on parylene C layers. Perfusion holes can help minimizing the excessive cerebrospinal fluid (CSF) around the electrode sites (*53*). The Ti hard mask and protection layer was removed by BOE and rinsed with DI water, leaving fresh Au surface at the electrode sites. The parylene C device was released from the Si carrier wafer by removing Micro-90 with DI water. The bonding pad region of the probe (Cr/Au) was bonded with ACF and commercial off-the-shelf ribbon cables and additionally encapsulated by kapton tape to prevent saline or CSF leakage (**fig. S10C**). To enhance recording qualities, PEDOT:PSS was electrodeposited on the surface of Au electrode sites using the technique described above. The surface probe consists of two column 32 channels along 3.12 mm in a laminar manner and the diameter and pitch of electrode sites is 20 and 100 µm, respectively (**fig. S10, D to F**).

Electrochemical impedance at 1 kHz was measured using Intan RHD2000 USB interface board (Intan Technologies) in 1X phosphate buffer saline (PBS) solution with a stainless-steel needle as a reference electrode (**fig. S10G**). The average impedance magnitude of Au electrodes was 739 ±

85.4 kilohms, which was decreased to 33.6 ± 3.21 kilohms after PEDOT:PSS electrodeposition on Au electrodes, with uniform distribution of the impedance across the channels (**fig. S10H**).

### 3. Mechanical characterizations of μSEEG electrodes

#### 3.1. μSEEG insertion benchtop tests

Insertion of the prototype μSEEG was tested on a phantom brain model. A transparent brain phantom gelatin model was prepared by mixing and dissolving a gelatin powder (Knox; Kraft Foods, Inc.) with weight concentration of 5.3 %. Knox gelatin with this concentration was reported to provide similar shear modulus to the mouse brain (*54*) and its transparency helps in visualization of the μSEEG electrode movement in the gel. We designed and printed a custom-made 3D printed holder that can be attached to the stereotaxic micromanipulator while holding the prototype μSEEG (**fig. S5A**). The prototype μSEEG, held by the 3D print holder, was slowly inserted into the gelatin using the z-axis control handle of the stereotaxic micromanipulator. After the prototype μSEEG was inserted, the stainless-steel stylet was manually grasped by a tweezer and extracted from the prototype μSEEG. The implantation of the prototype μSEEG and displacement after stylet extraction were recorded using a camera with a macro lens (**fig. S5B**). The prototype μSEEG was implanted without any deformation with help of the stylet. Also, the displacement of the prototype μSEEG before and after stylet extraction was less than 10 μm.

#### 3.2 Pull measurement of μSEEG

We investigated tensile stress of μSEEG using pull measurement to exam mechanical stability of our electrode and compare it with clinical depth electrode (PMT electrode). The μSEEG was integrated on the pressure sensor system (voice coil-powered linear actuator system with internally integrated force sensor; V-275 PIMag Voice Coil Linear Actuator). A customized 3D printed sample mount was attached onto the actuator (**fig. S4A**) and a tip of the μSEEG was placed on the sample mount followed by strong fixation using epoxy. Rest of the electrode was attached on a main body of the system and fixed by epoxy as well (**fig. S4B**). We applied force to the actuator to exam tensile strength upon the μSEEG break (indicated by red arrow) and obtained that the critical tensile strength of the μSEEG is 1 Mpa (16 mN) – 2.5 MPa (37 mN). Meanwhile, the PMT electrode that was anchored on two polyurethane tube regions around a Pt contact was damaged when 14 kPa (48 mN) was applied to the PMT electrode (**fig. S4C**).

#### 3.3 Mechanical stability of PtNR

PtNR μSEEG electrodes (N = 3) after extraction from the NHP brain was investigated by SEM (**fig. S8**). It was observed that PtNRs remained intact underneath the adsorbed tissue that are left on the surface of the electrodes (**fig. S8A**). A higher magnification SEM image (**fig. S8B**) shows PtNRs morphology similar to that of before implant (**Fig. 1B**).

### 4. Rat recording

#### 4.1. Acute rat recordings: surgical procedures and sensory stimulation

Acute *in vivo* electrophysiological recordings were performed on the rat primary somatosensory "barrel" cortex (S1) with the μECoG electrode and the μSEEG electrode. The rat was anesthetized and craniotomy was performed under isofluorane anesthesia. The body temperature of the rat was maintained at 37 °C with a heating pad. Craniotomy and dura removal were performed over the right barrel and surrounding cortical region. Following electrode placement, the rat was transitioned to ketamine/xylazine anesthesia for recording.

Tactile stimulation was performed by delivering air puffs to the whisker pad. Air puffs were pressure-injected through a glass micropipette using a PV830 pneumatic picopump (World Precision Instruments, Inc.) with 1 s pulses (n>10 trials per location). The contralateral (left) whiskers with respect to the recording sites were deflected by air puff (± 2 mm). First, the whole contralateral whiskers (multi-whisker) were stimulated. Then single whiskers (C1-3, D1-3, E1-4) were stimulated by placing the pipette as close as possible to each whisker to avoid deflection of the neighboring whiskers. Recording data were collected for 60 s for each whisker.

4.2. Acute rat recordings: histology of the rat brain

Prior to the insertion to the cortex, the back of the tip of the µSEEG electrode was painted with a fluorescent dye, DiI (1,1′-Dioctadecyl-3,3,3′,3′-tetramethylindocarbocyanine perchlorate, 0.1% in ethanol; Invitrogen) to visualize the µSEEG electrode track in histopathological sections and verify the placement of the µSEEG electrode with respect to cortical depth. Following the electrophysiological recordings, rats were sacrificed and perfused with 4% paraformaldehyde (PFA), and the brains were fixed for 2 hrs in 4% PFA. Brains were transferred to a 30% sucrose solution, and then embedded in OCT matrix and stored at -80. Slices were sectioned on a cryostat at 50 µm. Free-floating sections were washed in phosphate buffered saline (PBS) + 0.1% Triton for 10 min, and then incubated in 1:50 Neurotrace 500/525 (ThermoFisher) for 30 min. Sections were washed in PBS + 0.1% Triton for 10 min, followed by PBS. Sections were mounted on slides and coverslipped using ProLong Gold (ThermoFisher). Slices were imaged using a Keyence BZX-700 at the UCSD Department of Neuroscience Microscopy Core (supported by NS047101). From the histology result (**fig. S11**), the tip of the acute µSEEG electrode was estimated to be 1750 µm deep in the cortex, reaching layer VI.

4.3. Chronic rat recordings: implantation of µSEEG electrode, surgical procedures, and sensory stimulation

The short 32 channel µSEEG electrode for chronic rat recordings consists of 32 channels of microcontacts that are spread along the 2-mm-scale µSEEG electrode, which are connected to the metal pads for interfacing with PCB (**fig. S15A**). The metal pads and PCB are bonded by Ag epoxy. The back side of the PCB has electrical and mechanical contact with zero-insertion-force (ZIF) connector. The ZIF connector was activated with a lever to close the contact after flat flexible cable (FFC) insertion into the ZIF connector to make secure electrical and mechanical contact.
For implantation of the chronic µSEEG electrode, first, a C-shape mount was fixed onto the skull near the craniotomy using screws and it was then bonded with the headstage using a super glue. Then, the chronic µSEEG electrode with PCB and ZIF with FFC inserted was temporarily attached on the glass slide and the glass slide was connected to the micromanipulator (**fig. S15B**). During the implantation, position of the chronic µSEEG electrode was precisely controlled by micromanipulator. After the implantation, FFC was unplugged from the ZIF and the chronic µSEEG electrode with PCB and ZIF are securely protected by the headstage and fixed onto the skull due to the screw-fixed mount. The craniotomy was sealed by ultraviolet curable medical glue (Tetric EvoFlow). Stimuli to induce neural activity included whisker, paw, and trunk mechanical stimulation including air puffs delivered >10 times/trials and the recording data from each trial were averaged.

4.4. Chronic rat recordings: histology of the rat brain

At the termination of the chronic recordings (by four weeks after implant), brains were removed and briefly (2 hrs) fixed in 4% PFA. The cortices were removed and flattened between glass slides in PBS, and then the flattened cortex further fixed in 4% PFA overnight, and then allowed to sink in 30% sucrose. For measuring implantation scar, rats were implanted with a dummy clinical electrode or µSEEG and allowed to recover for two weeks (n = 1 / electrode). Animals were perfused with 4% PFA and then sunk in 30% sucrose. Whole brains or flattened cortices were sectioned on a cryostat in the horizontal plane from at 40 µm. Sections were washed with PBS and stained overnight with one or more of the following: chicken anti-NeuN (1:500, Sigma ABN91), Cy3-conjugated mouse anti-GFAP (1:1000, Sigma C9205), rabbit anti-VGLUT2 (1:500, Abcam ab216463). After wash, slices were incubated in secondaries of goat anti-chicken 488 (Sigma SAB4600039), goat anti-chicken 647 (Sigma SAB2600184), or goat anti-rabbit 488 (ThermoFisher A-11008) at 1:300 for 1 hour, rinsed, and mounted with ProLong with DAPI. Slides were imaged on a Nikon Eclipse Ti2-E with a DS-Q12 CMOS camera at the UCSD Nikon Imaging Core Images were analyzed using ImageJ (**fig. S7 A to D**). In our observations from one chronic (2 weeks) placement, immunochemistry revealed NeuN-positive cells in close (< 20 µm) proximity to the µSEEG electrode and lower GFAP intensity and spread (**fig. S7E** and **F**).

4.5. Rat recording methods

We successfully gathered depth recordings of all intended stimulus-evoked responses and gathered tissue for histological evaluation. These data were not blinded, and we did not include a randomization of subject selection in this study. The data from the µSEEG electrode was acquired at 15 – 20 kHz and filtered by default Intan settings with cutoffs of 1 Hz to 7.5 kHz. The impedance tests of the electrodes during the experiments and recordings in the animals carried out using the Intan RHD2000 software from Intan Technologies (Los Angeles, CA). Data was extracted and processed using MATLAB (Mathworks, Natick, MA).

### 5. Pig recording

5.1. Pig surgical details and spinal cord stimulation

The pig experiments were approved by the University of California San Diego (UCSD) Institutional Animal Care and Use Committee. Yucatan pigs were induced with isoflourance and intubated. Once anesthetized, the animal was mounted into a stereotaxic frame in the prone position. Vital signs were monitored including heart rate, blood pressure, EtCO2, respiratory rate, and blood oxygenation. The cranial surgical site was centered over the motor cortex of the frontal lobe and somatosensory cortex of the parietal lobe. After immobilization, a skin incision (2-4 inches long) was made and unilateral craniotomy was performed using a high speed surgical drill. The bone was removed to create a window approximately 25 mm x 15 mm. The underlying dura was incised and reflected medially towards the sagittal sinus. The cortex was hydrated with normal saline solution. A flexible sterile multielectrode implant was then placed on the surface of the exposed cortex. After placement of the electrode into the cortical surface, the electrode ribbon (<1 cm in width) was connected to an Intan recording controller. For spinal exposure, a midline incision and laminectomy were performed with a Kerrison rongeur to sufficiently expose the target thoracolumbar enlargement of the spinal cord. Following laminectomy, a longitudinal incision was made in the dura and arachnoid tissue was dissected to allow for exposure of the spinal cord. Spinal cord stimulation was performed with a handheld Ojemann stimulator with two ball tips (Radionics Inc., Burlington, MA) with 0.5 cm spacing between electrode tips. Isoflurane anesthesia was halted following completion of surgical procedures and was replaced with IV Propofol for the duration of the stimulation testing.

5.2. Pig recording methods

We successfully gathered depth recordings of all intended stimulus-evoked responses. These data were not blinded, and we did not include a randomization of subject selection in this study. The data from the µSEEG electrode was acquired at 15 – 20 kHz and filtered by default Intan settings with cutoffs of 1 Hz to 7.5 kHz. The impedance tests of the electrodes during the experiments and recordings in the animals carried out using the Intan RHD2000 software from Intan Technologies (Los Angeles, CA). Data was extracted and processed using MATLAB (Mathworks, Natick, MA).

### 6. NHP recording

Experimental procedures on rhesus macaques were carried out in accordance with the Guide to the Care and Use of Laboratory Animals. All efforts were made to minimize discomfort, and the Institutional Animal Care and Use Committee at the Massachusetts General Hospital monitored care and approved all procedures. We tested µSEEG electrodes in two settings: 1) recording from a short µSEEG in an anesthetized NHP the operating room for recording visual cortical dynamics and 2) recording from a long µSEEG in an awake NHP with the electrode lowered through an implanted chamber with a standard Microdrive (**table S1**).

An intraoperative, intracranial neurophysiology recording were acquired from one adult male rhesus macaque (*Macaca mulatta*, age 11). The macaque was placed under general endotracheal anesthesia (isoflurane) and placed into a stereotactic frame (Kopf; Kujunga, CA). A craniotomy over the visual cortex was performed using standard anatomic landmarks, and cortex was carefully exposed and the short µSEEG electrode was implanted. Signals were recorded using a custom Intan Recording System as described previously (*29, 55*).

In the awake NHP preparation, one adult male rhesus macaque (*Macaca mulatta*, age 14) was implanted with two recording chambers that allowed access parietal lobe, temporal lobe, and thalamic brain regions. A fitted Microdrive was placed on the top of the chamber to allow insertion through the dura. Following recording, trajectories were calculated by mapping the depths calculated from the Microdrive to a preoperative magnetic resonance imaging (MRI) and a postoperative CT of the chamber locations using MANGO (San Antonio, TX). To visualize the locations in 3D, we used the Scalable Brain Atlas with the Calabrese atlas (https://scalablebrainatlas.incf.org/; (*46, 47*) exported into Blender (https://www.blender.org/). The electrode was lowered acutely to three different depths to record neural activity. No clear adverse behavioral effects were observed before or after electrode implant or explant. Recordings of the long µSEEG involved the use of the 1024 Intan Recording system with a 1024 channel Intan chip clamped device developed for recording thin film microelectrodes (*30*).

For both preparations, data from the µSEEG electrode was acquired at 30 kHz and filtered by default Intan settings with cutoffs of 1 Hz to 7.5 kHz. The majority of the data was acquired using OpenEphys acquisition graphic-user interface software ((*56*); http://www.open-ephys.org/), with the impedance tests of the electrodes during the experiments carried out using the Intan RHD2000 software from Intan Technologies (Los Angeles, CA). Data was extracted and processed using

MATLAB (Mathworks, Natick, MA). Data recorded from the µSEEG electrode was down-sampled to 1000 Hz.

### 7. Human recording

7.1. Recordings in human cortex the operating room

Intraoperative recordings were performed in two participants who underwent a neurosurgical procedure at Massachusetts General Hospital (MGH). This study was approved by the Partners Institutional Review Board (now the Mass General Brigham Institutional Review Board which covers MGH). Both participants were involved voluntarily, provided informed consent, and were informed that participation in the experiment would not alter their clinical treatment in any way and that they could withdraw at any time without altering their clinical care. The patients were approached following already being scheduled to remove a portion of the left anterior temporal lobe for the treatment of epilepsy or tumor removal. The possibility of conducting research recordings were only discussed with each patient after the decision to proceed with the surgery had been made. Neither patient was medically unstable or required emergency or urgent surgery. All decisions to proceed were made following consultation with the treating neurosurgeon and clinical team. Further, we did not enroll patients with clearly impaired decision-making abilities (as determined by the primary clinical team or physician caring for the patient). The time allowed for dedicated research recording was limited per subject to minimize risk. We tested the short µSEEG electrodes, which were inserted into tissue the clinical team identified would be resected following recording. One participant (HS1) was under general anesthesia but underwent clinically indicated stimulation of the cortical surface prior to resection. The second participant (HS2) underwent awake cognitive testing under monitored anesthesia care (MAC) with electrocorticography (intracranial EEG) during surgery.

Neural signals were recorded using a custom Intan Recording System as described previously (*34, 55, 57*). Data from the µSEEG electrodes were acquired at 30 kHz and filtered by default Intan settings with cutoffs of 1 Hz to 7.5 kHz using the OpenEphys acquisition graphic-user interface software ((*56*) http://www.open-ephys.org/), with the impedance tests using the Intan RHD2000 software from Intan Technologies (Los Angeles, CA).

7.2 Auditory Task in the Operating Room

The Auditory M-sequence block involves 100 trials where each trial involves a red fixation cross on a grey background while one of either two tones are played in the background. The choice of tone, whether 250 Hz or 500 Hz, per trial was randomly sampled from sequences of black or white designations pulled from an m-sequence distribution (*56, 58, 59*). These frequencies and tones have been used before to demonstrate auditory specificity intracranially in humans (*60*).

### 8. Neural Data Analysis

Data was extracted and processed using MATLAB R2020a (Mathworks, Natick, MA). Data recorded from the µSEEG electrode was down-sampled to 1000 Hz. To remove noise, the data were re-referenced by subtracting the median signal of all the channels. Local field potentials (LFPs), multi-unit activity power (300-2000 Hz) high gamma activity (HGA; 65-200 Hz) evoked by tactile stimuli were investigated. Power spectral analyses to examine the dynamics in the different conditions were performed by taking the real value of the Morlet wavelet coefficient (power) at a 1-Hz spectral resolution using a moving window of 0.5 sec moving every 10 ms, calculated using the Fieldtrip toolbox (www.ru.nl/fcdonders/fieldtrip, (*61*)). In the case of direct electrical stimulation, stimulation artifact was removed using a Tukey-windowed median filter in the 20 ms around the onset of the pulse for the single pulses (*62*). Other measures of multi-unit activity involved measuring high gamma power (70-190 Hz) by taking the absolute value of a Hilbert transform following band pass filtering.

Current source density (CSD) provides information on the distribution of synaptic inputs for the generators (*63*). Here, CSD was calculated from LFP using the δ-source inverse CSD (iCSD) method, refined by Petterson et al. (*64*):

$$C = F^{-1}\Phi$$

where $C$ and $\Phi$ represents matrix for CSD and LFP, respectively. The transformation matrix $F$ is described by

$$F_{ji} = \frac{h^2}{2\sigma}\left(\sqrt{(j-i)^2 + (R/h)^2} - |j-i|\right)$$

where $h$ is the spacing between adjacent electrode, $\sigma$ is the conductivity of the cortex and $R$ is the radius of infinitely thin current-source discs. Here we set the $\sigma$ and $R$ as 0.3 S/m and 0.25 mm, respectively. Since CSD is calculated from LFP data of adjacent electrodes, each column was separated for calculation of CSD and the column that included a greater number of low impedance (<500 kΩ) channels (30 out of 32) was selected for the CSD analysis.

We detected bursts and calculated the burst suppression ratio (BSR) using an automated method (*44*) (https://github.com/drasros/bs_detector_icueeg). On the per-channel basis, this method labels each time sample as either burst or suppression. Briefly, the method uses the previous data with each channel and applies the following equations:

μt = β μt-1 + (1-β) xt

σt2 = β σt-12 + (1-β) (xt - μt)2

zt = δ[σt2 < θ]

Where xt is the value of the normalized signal of one channel at time t, μt and σt2 are current values of the recursively estimated local mean and variance, respectively. Finally, zt is an indicator function that labels each data point as either a burst (0) or suppression (1). The value of β determines the balance between the effect of recent and past data set based on previously trained data (*44*). The classification threshold θ (i.e., the value above which a data point should be classified as burst) was adjusted to evaluate our dataset visually with values of θ = 100, 2000, 4000. The value of θ = 2000 was selected to reliably identify burst and suppression induced by general anesthesia. The burst suppression ratio for each recording in an anesthetized patient (N = 1) and NHP (N = 1) was evaluated as the proportion of suppression-labeled samples in a moving window (1 s duration, no overlap).

Spike sorting was performed on the long μSEEG recordings from an NHP using Kilosort 2.5 (*45*); https://github.com/MouseLand/Kilosort) with further determination of single units versus multi-unit activity performed in post processing using Phy (https://github.com/cortex-lab/phy ) and then manually curated using in-house MATLAB code to visually inspect the template as well as the waveforms assigned to each cluster. The Kilosort 2.5 parameters included: Nblocks = 0 – as no additional registration was needed according to spiking activity after the manual registration; Threshold [10, 11]. Clusters were separated into single units and MUA. Clusters of waveforms were classified as MUA if there was a mixture of distinct waveforms (examined in Phy) as well as a complicated (and abnormal) autocorrelogram. For all clusters, we measured the spike duration, peak-trough ratio, and amplitude measures (**Fig. 4**; code adapted from (*65*); https://github.com/jiaxx/waveform_classification).

### 9. Statistical Analysis

All statistical comparisons were performed using non-parametric measures, so we did not test for normality. We tested comparisons with the Kruskal–Wallis test for non-equivalence of multiple medians followed by the use of the *post hoc* Tukey-Kramer method to determine statistically separable groups. We used the Wilcoxon rank-sum test (two-sided) for comparisons between individual medians. We tested if values were significantly different from zero using the Wilcoxon signed rank test. We corrected by adjusting the target p-value (0.05) with a Bonferroni correction for the number of comparisons being done.

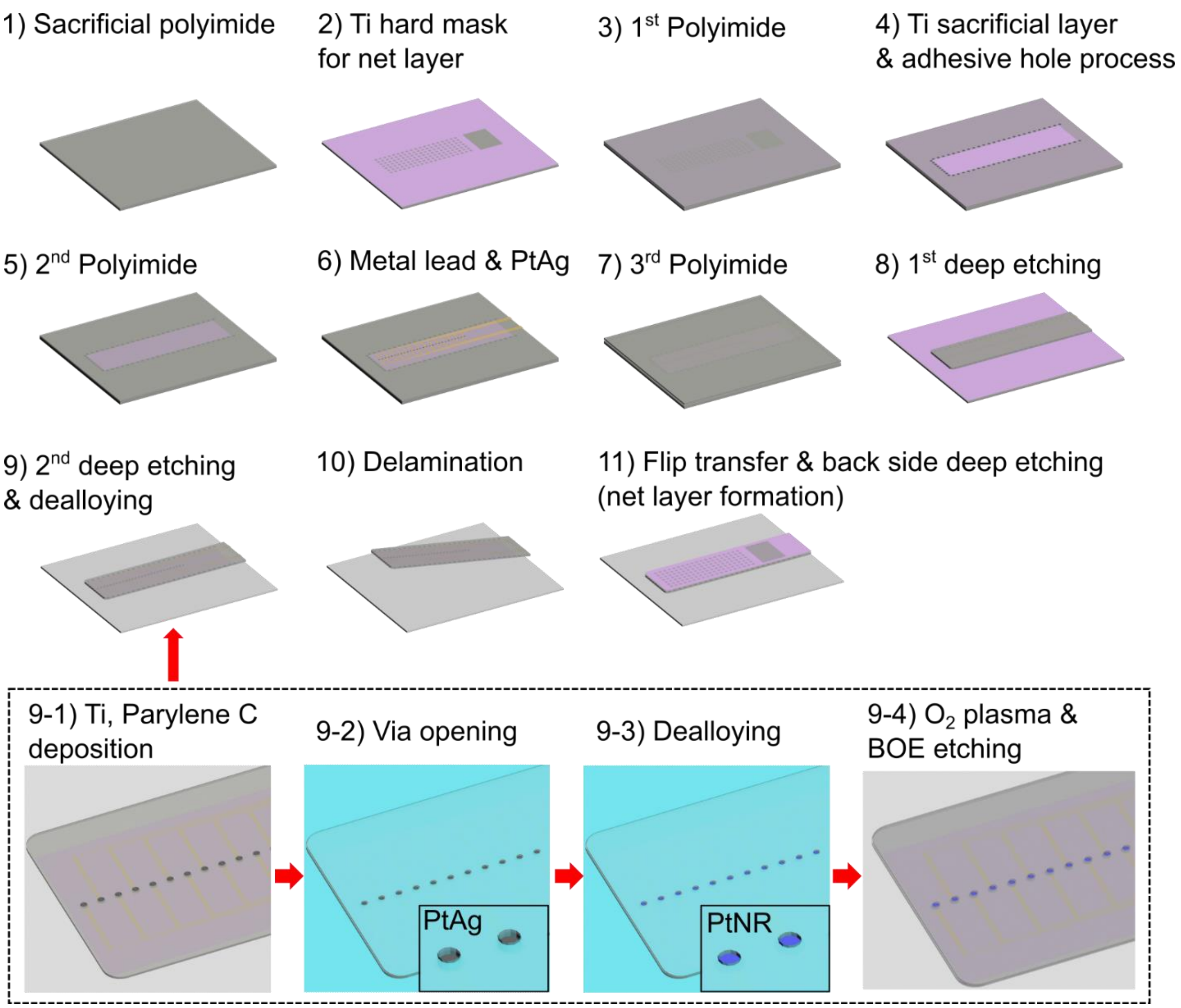


**Fig. S1**. **Fabrication process of μSEEG electrode**.

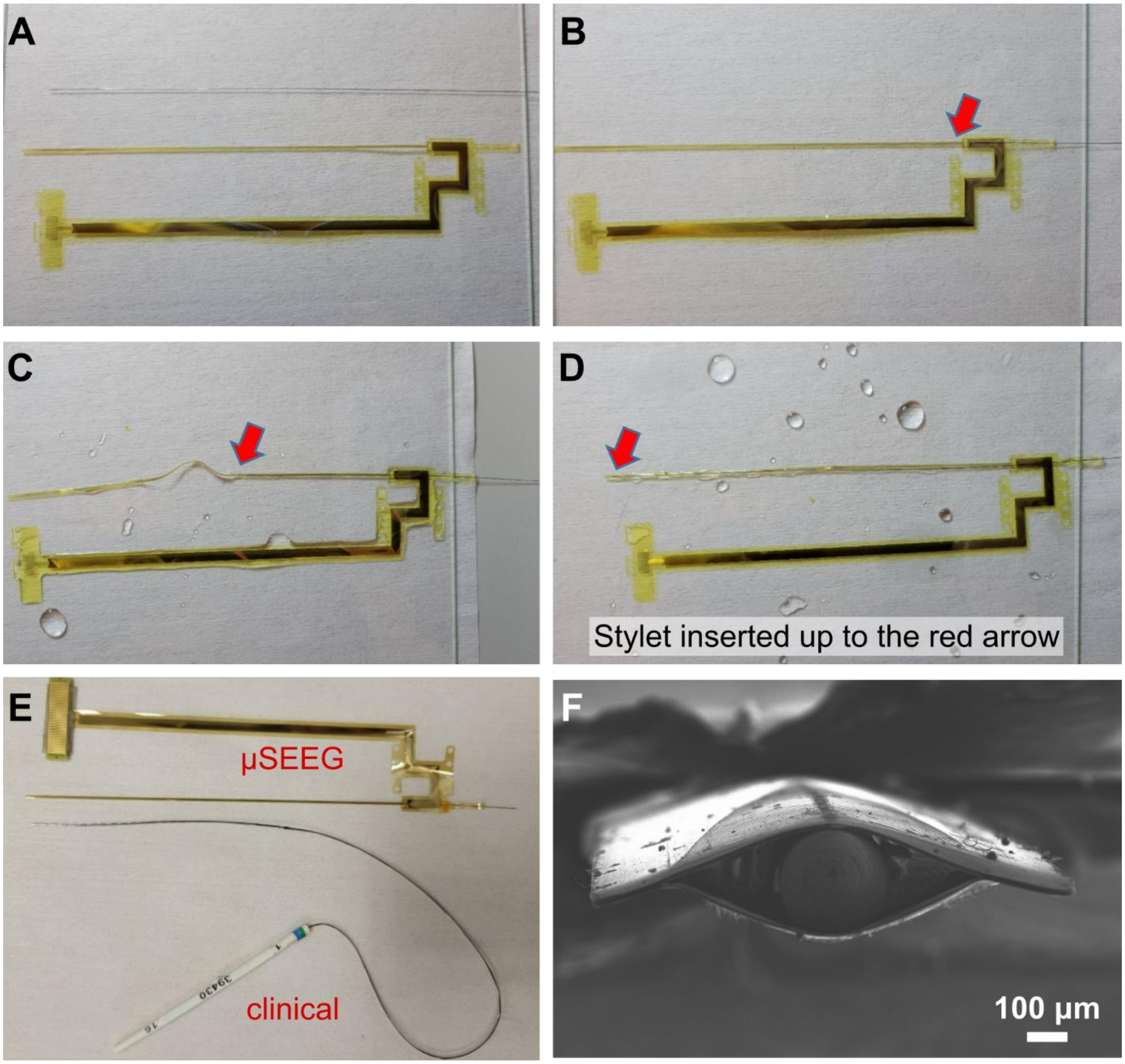


**Fig. S2**. **Stylet insertion into the μSEEG electrode**. (**A**) Before insertion. (**B**) Stylet aligned to the sheath. (**C**) During insertion. (**D**) Stylet insertion all the way to the tip. (**E**) Scale comparison with conventional SEEG. (**F**) Cross-sectional SEM image of the μSEEG.

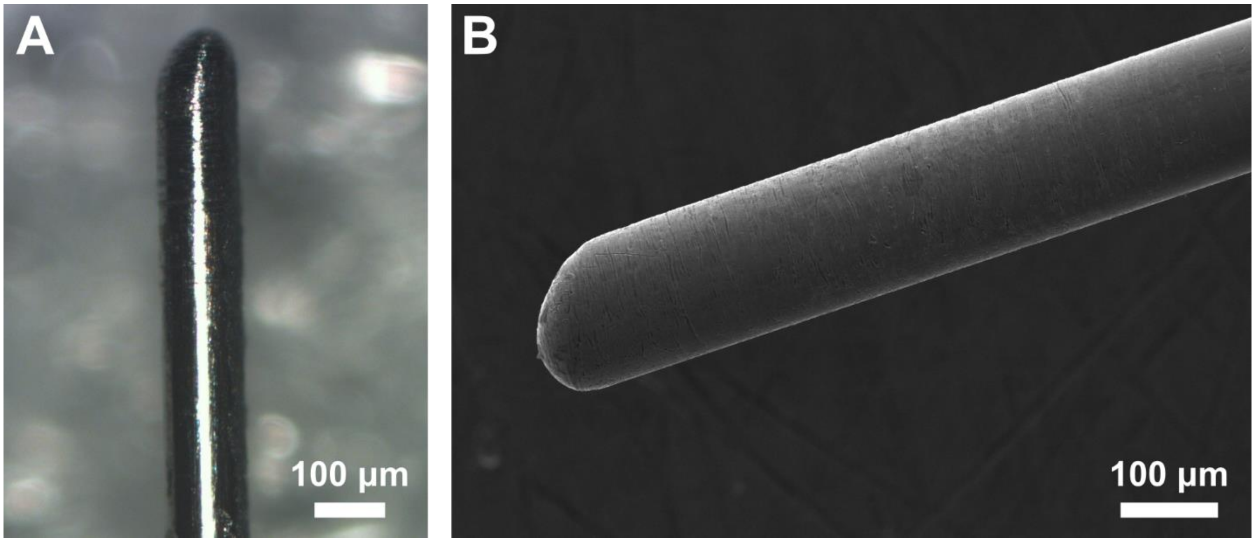

**Fig. S3**. **Medical grade stainless-steel stylet tip.** (**A**) OM image and (**B**) SEM image of a polished stainless-steel stylet.

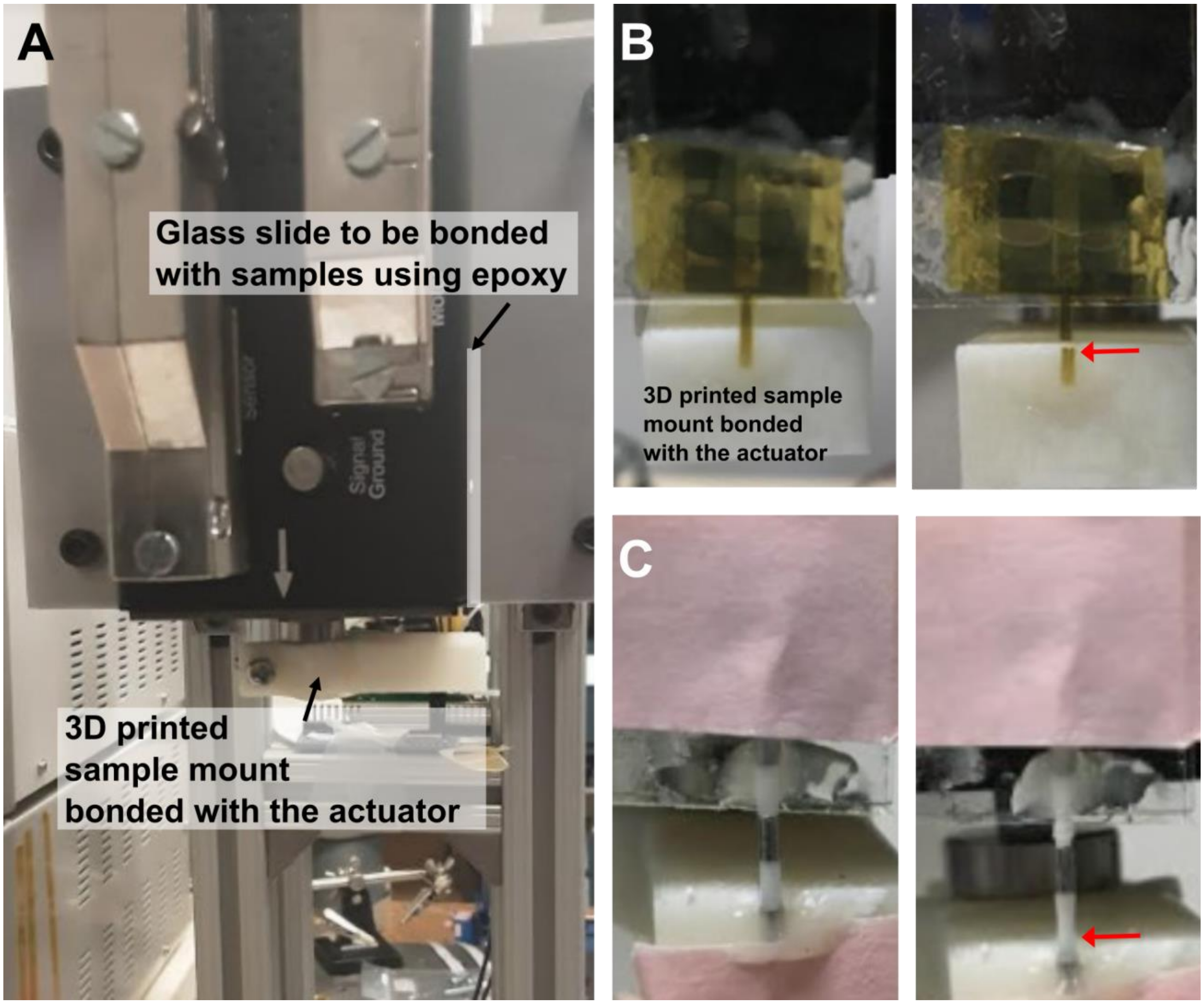

**Fig. S4. Pull measurement setup and result.** (**A**) Measurement setup. (**B**) Our μSEEG and (**C**) 1.2 mm diameter PMT depth electrode integrated on the measurement setup.

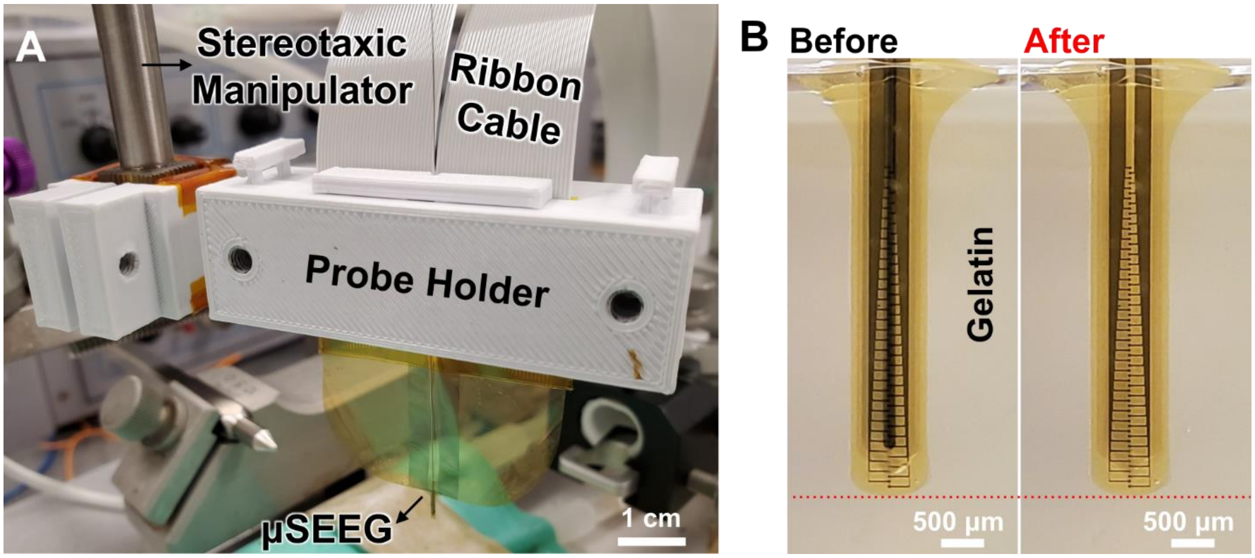


**Fig. S5. Acute insertion into a phantom brain model.** (**A**) An optical image of the 3D printed holder and the µSEEG electrode attached to the stereotaxic manipulator. Scale bar is 1 cm. (**B**) An optical image of the µSEEG electrode tip inserted in the gelatin model, before and after stylet extraction. Scale bar is 500 µm.

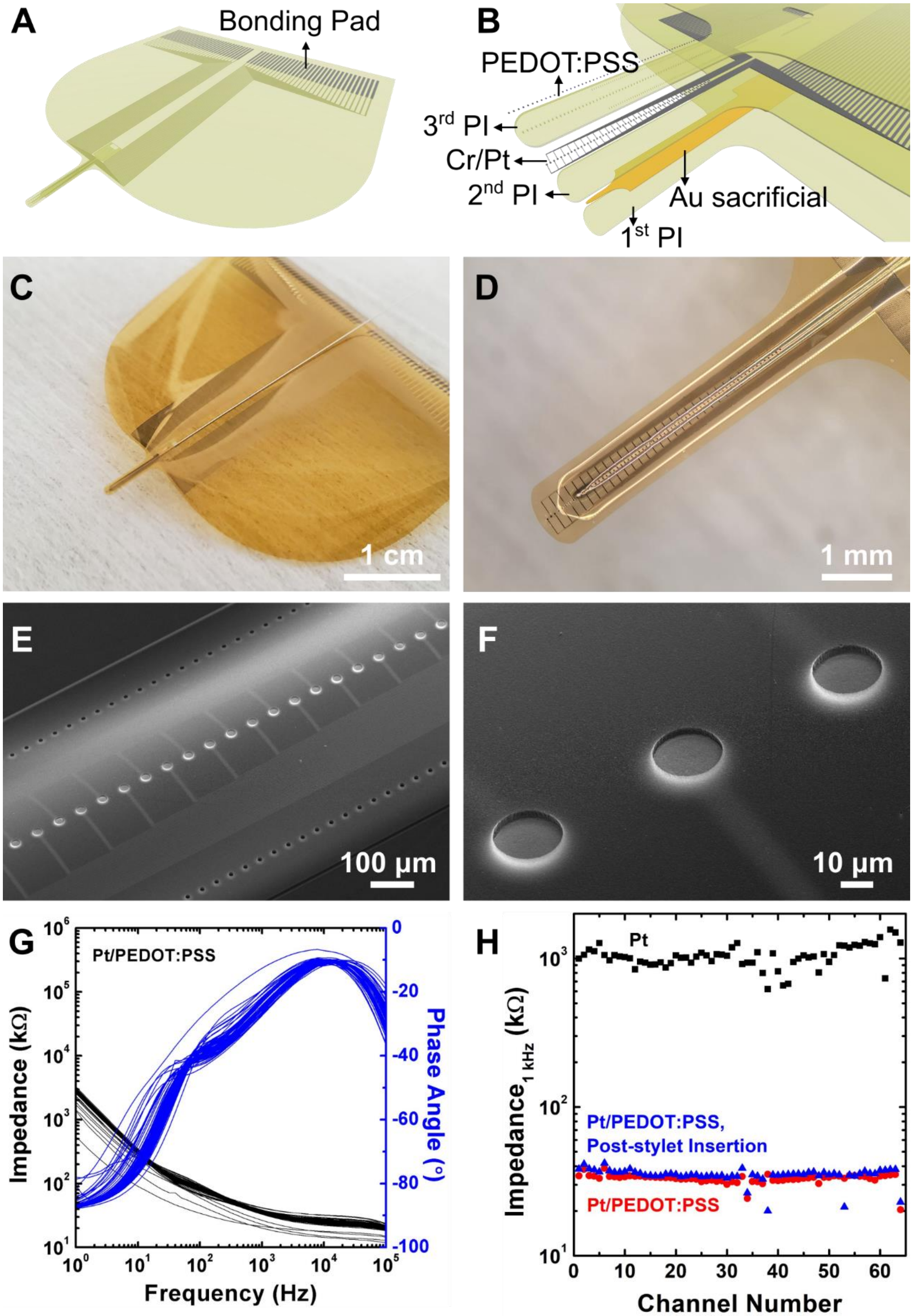


**Fig. S6. Manufacture and validation of the short PEDOT:PSS µSEEG in polyimide (PI)**. (**A**) A schematic illustration of a prototype 64 channel µSEEG electrode layout. (**B**) Exploded view of the layout at the electrode sites region. (**C**), (**D**) Optical images of the µSEEG electrode after stylet insertion. (**E**), (**F**) SEM image of μSEEG electrode showing PEDOT:PSS on Cr/Pt after stylet insertion. (**G**) Electrochemical impedance spectra of Pt/PEDOT:PSS recording electrodes. (**H**) Electrochemical impedance magnitude at 1 kHz of each channel before and after stylet insertion and comparison with Pt.

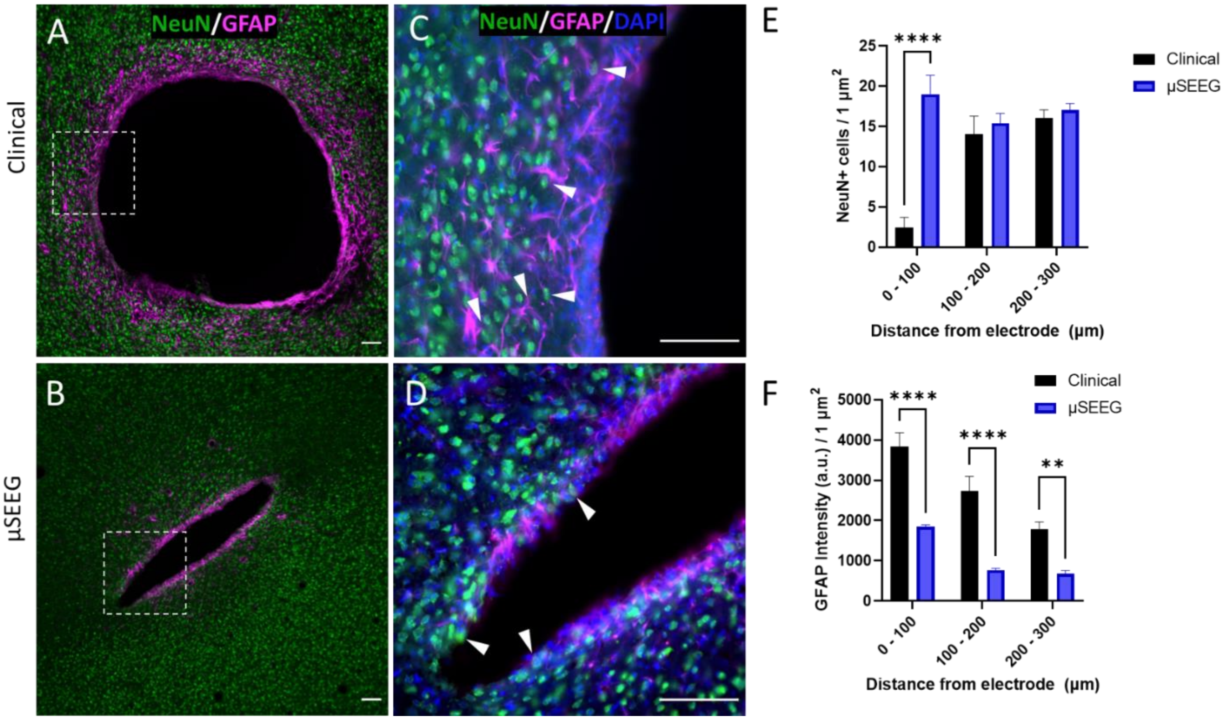


**Fig. S7. GFAP and NeuN reactivity folloing two weeks of electrode implantation.** Wide-field view of horizontal cortical section following (**A**) clinical or (**B**) µSEEG electrode implantation. NeuN is pseudocolored green, GFAP magenta, and DAPI blue. (**C**) and (**D**) are magnifications of the boxed areas in (**A**) and (**B**) to show NeuN reactivity adjacent to electrode placement. Represnetative NeuN-positive cells are indicated by arrows. (**E**) Average number of NeuN+ cells per 100 µm2 sampled randomly at 0-100, 100-200, and 200-300 µm from the electrode. (**F**) Average GFAP intensity of 100 $\mu m^2$ boxes sampled randomly at 0-100, 100-200, and 200-300 100 µm from the electrode. Data are plotted as mean ± SEM, **** $P < 0.0001$, ** $P < 0.01$ by 2-way ANOVA followed by Sidak's multiple comparisons test. Scale bar is 100 µm.

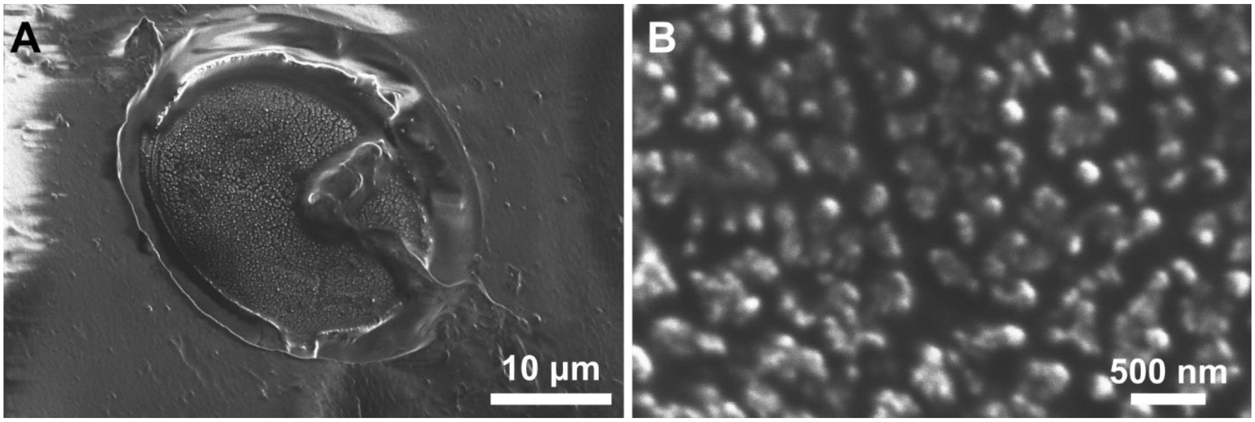


**Fig. S8. SEM images of μSEEG electrodes upon extraction.** (**A**) Low magnification image showing whole single PtNRs contact and biological tissue and (**B**) High magnification image of PtNRs contact.

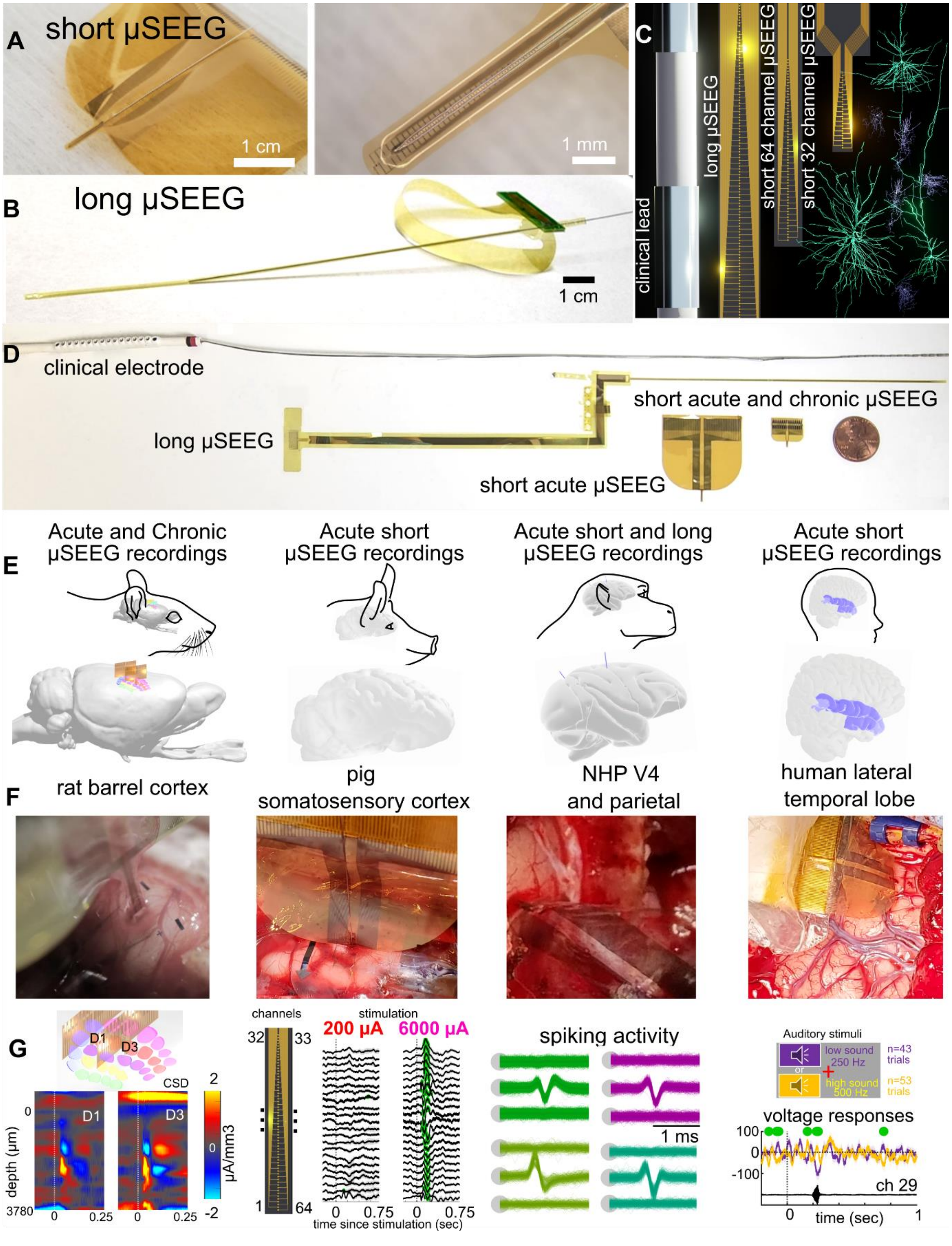


**Fig. S9. µSEEG Electrodes for Recording Cortical Neurophysiological Activity in Multiple Species.** (**A**). The short µSEEG with 64 microelectrode contacts at 60 µm contact to contact spatial resolution. (**B**) The long µSEEG with 128 microelectrode contacts at 60 µm contact to contact spatial resolution. (**C-D**) Diagram and photo of the relative scale of human cortical neurons relative to a clinical SEEG lead and a short µSEEG electrode (*35-37*). (**E**) Different species and brains along with the locations of the electrodes for neural recordings in this study. (**F-G**) Images of the different preparations inserted into cortex with example neurophysiological recordings below the images.

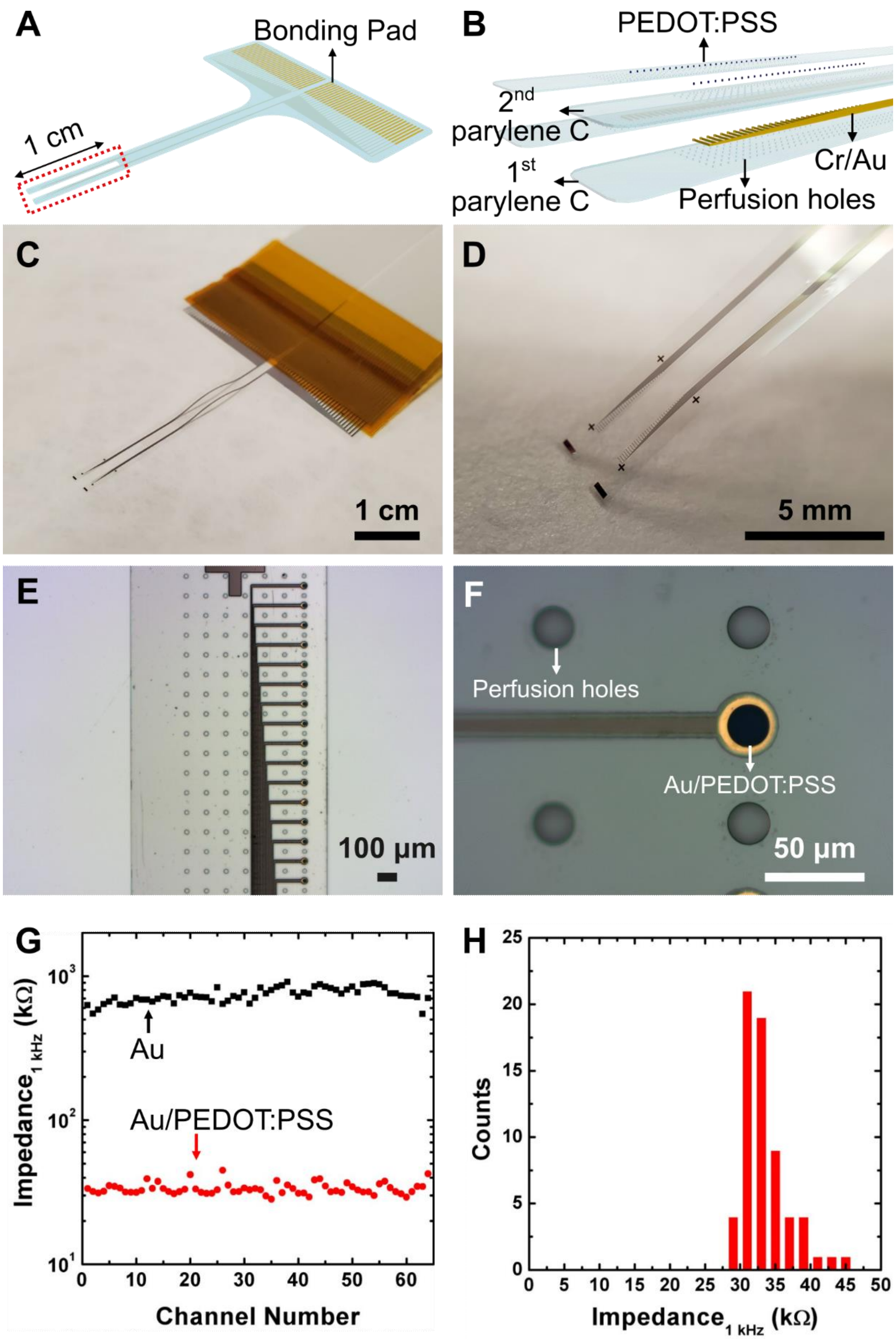


**Fig. S10. Manufacture and characterization of the μECoG for comparing surface to depth electrode capabilities.** (**A**) A schematic illustration of μECoG electrode layout. Red dotted square indicates electrode sites region. (**B**) Exploded view of the layout at the electrode sites region. (**C**), (**D**) Optical images of the μECoG electrode. Scale bar is (**C**) 1 cm and (**D**) 5 mm. (**E**) Top-view optical microscopic image of the electrode sites region. (**F**) Magnified top-view optical microscopic image showing PEDOT:PSS on Cr/Au electrode sites and perforation holes. (**G**) Electrochemical impedance magnitude at 1 kHz of each channel of μECoG electrode before and after PEDOT:PSS electrodeposition. (**H**) Electrochemical impedance magnitude at 1 kHz histogram of Au/PEDOT:PSS.

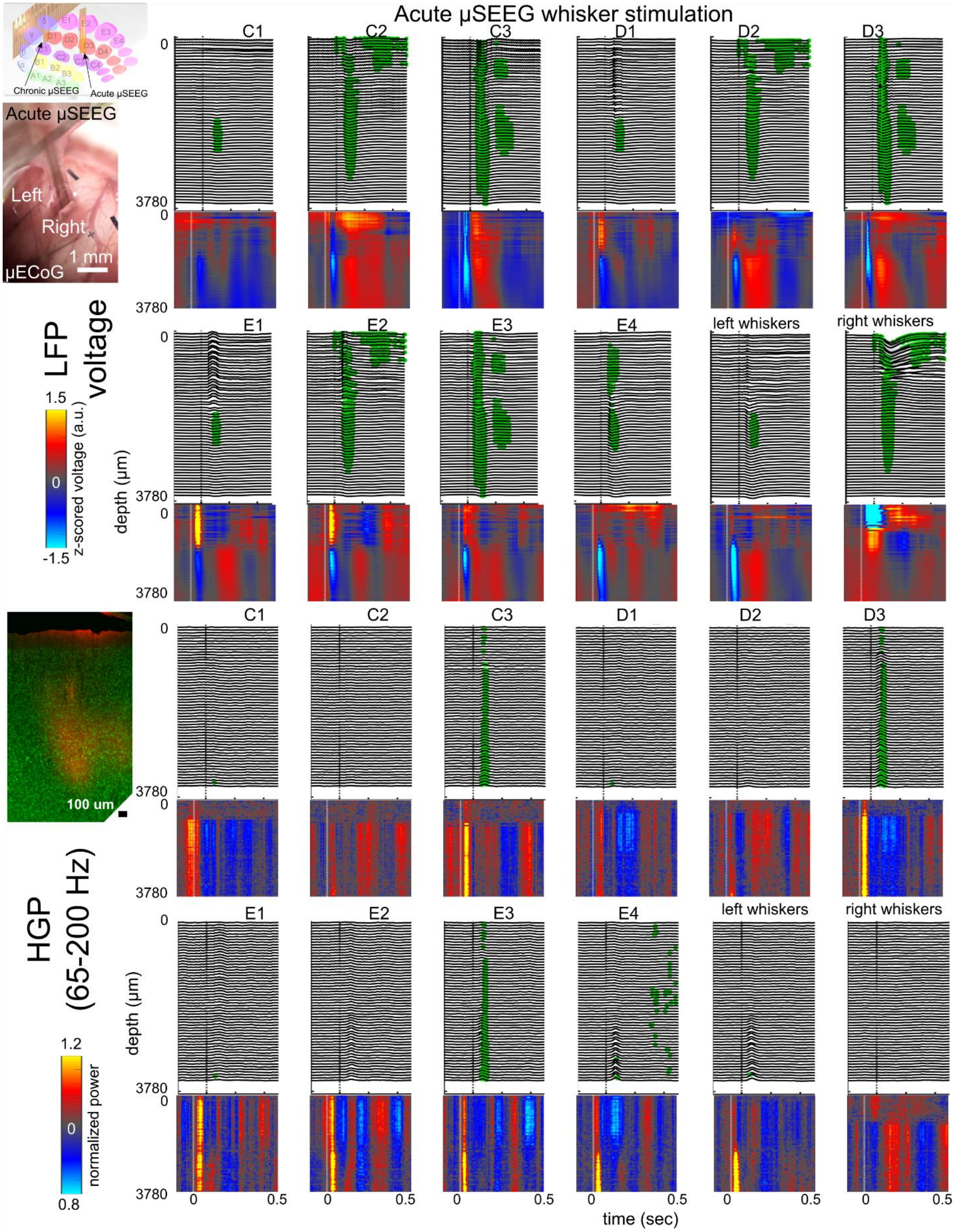

Acute μSEEG whisker stimulation
Chronic μSEEG
Acute μSEEG
Acute μSEEG
Left
Right
1 mm
μECoG
C1
C2
C3
D1
D2
D3
E1
E2
E3
E4
left whiskers
right whiskers
LFP voltage
z-scored voltage (a.u.)
1.5
0
-1.5
depth (μm)
0
3780
100 um
HGP (65-200 Hz)
normalized power
1.2
0
0.8
time (sec)
0.5

**Fig. S11. Acute laminar recording in the rat barrel cortex with the µSEEG with sensory stimulation.** Upper left: Location and 3D reconstruction of possible locations of the acute and chronic implantation of µSEEG electrodes for recording from the rat barrel cortex and images of the implanted µECoG electrode (left) and the µSEEG electrode (right). First, the µECoG electrode was placed on the cortical surface. At the gap between two columns of the µECoG electrode, the µSEEG electrode was implanted, and the stylet was removed using the previously described technique. Note some contacts are outside brain tissue on the µSEEG electrode. Voltage responses across the µSEEG electrode to whisker stimulation at different whisker locations for the voltages and high gamma power (HGP). Green dots indicate significantly different from 0.5 sec before (which is baseline) air puff stimulation to the whisker (Wilcoxon rank-sum test) per channel and across trials. Number of trials >10.

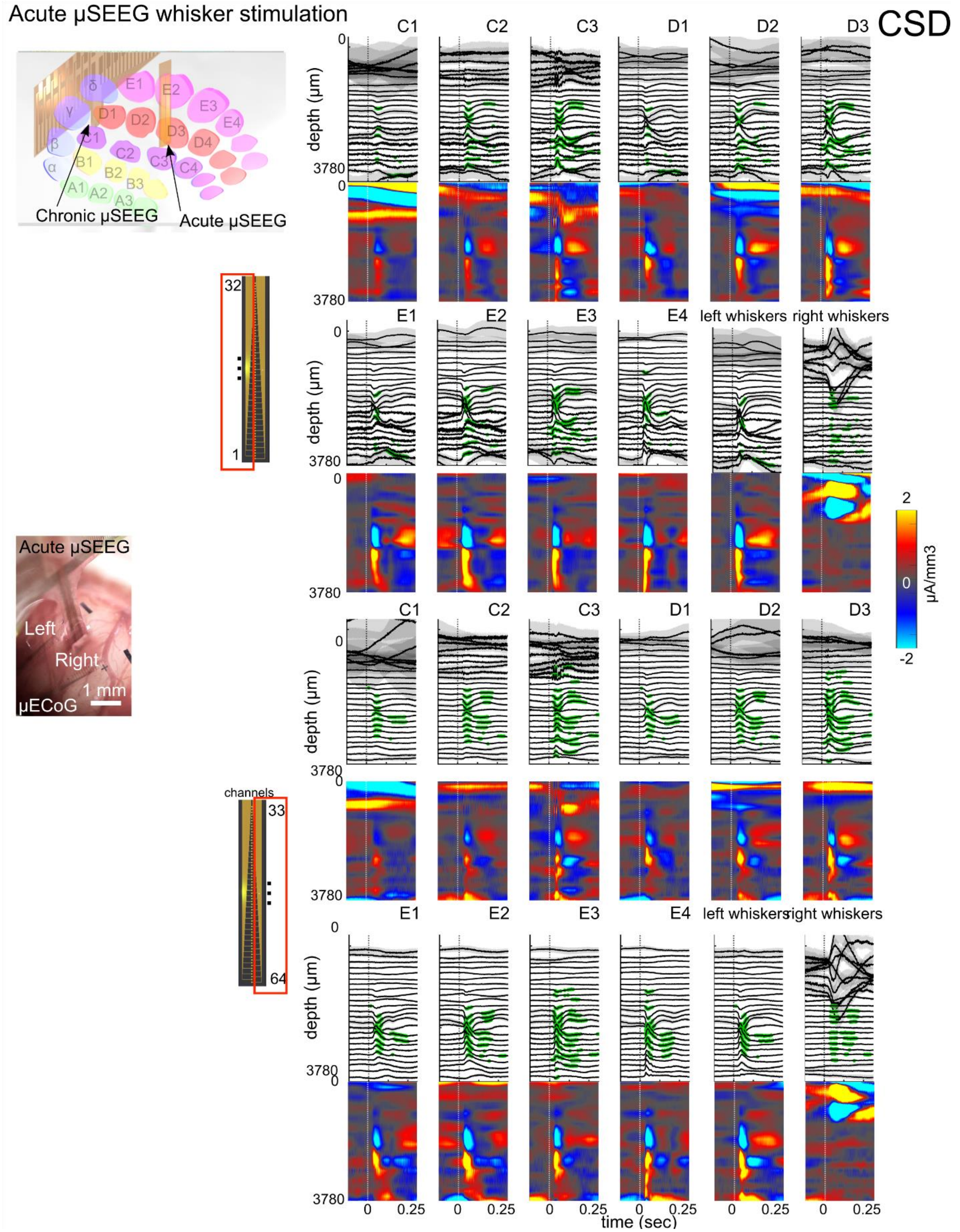


**Fig. S12. Acute laminar recording in the rat barrel cortex with the µSEEG with sensory stimulation.** Upper left: Location and 3D reconstruction of possible locations of the acute and chronic implantation of µSEEG electrodes d for recording from the rat barrel cortex and images of the implanted surface µECoG electrode and the µSEEG electrode. Note some contacts are outside brain tissue on the μSEEG electrode. Current source density (CSD) responses across the μSEEG electrode to whisker stimulation at different whisker locations. Green dots indicate significantly different from 0.5 sec before (which is baseline) air puff stimulation to the whisker (Wilcoxon rank-sum test) per channel and across trials. Number of trials >10.

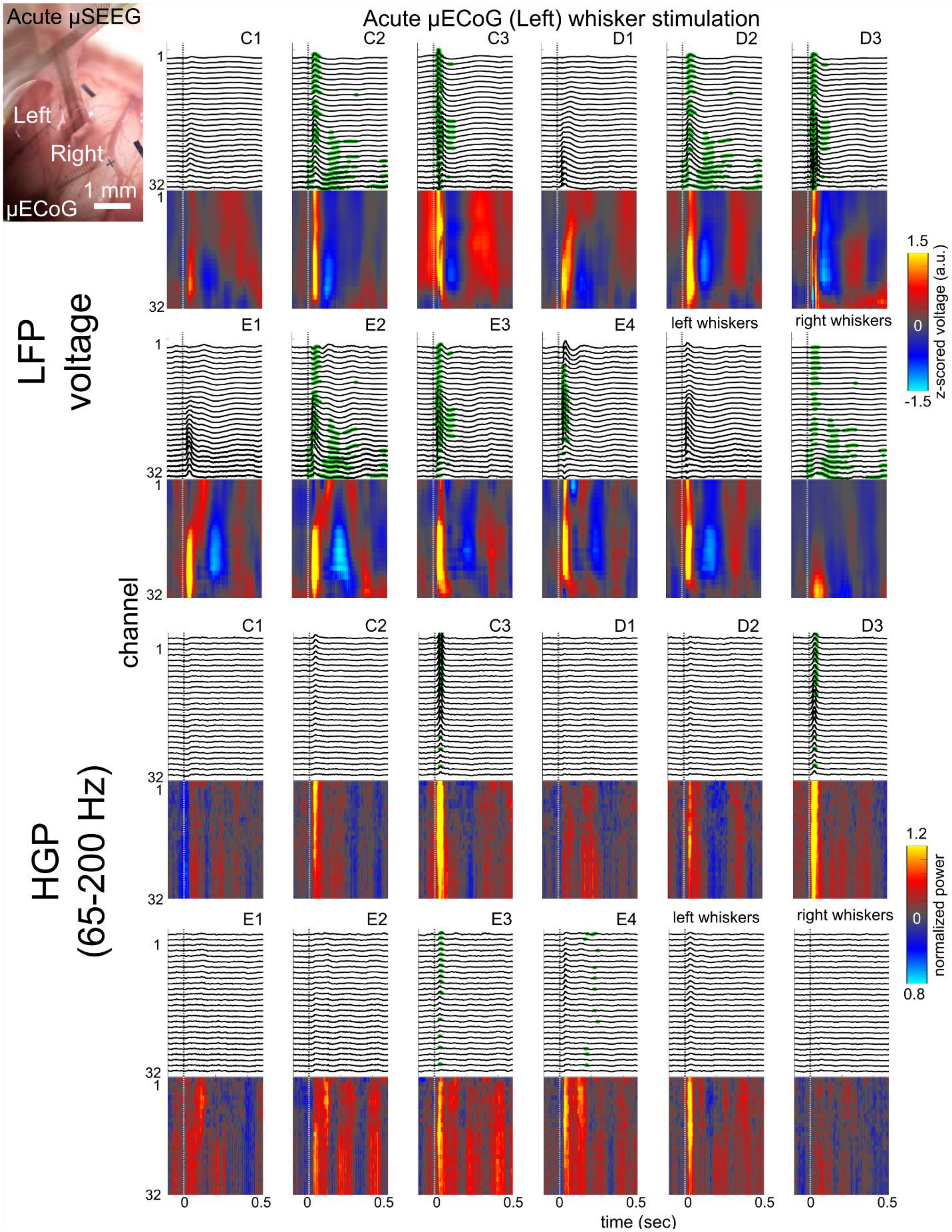


**Fig. S13. Acute surface recording with a strip of the µECoG over the rat barrel cortex with sensory stimulation.** Upper left: Location and 3D reconstruction of possible locations of the acute and chronic implantation of µSEEG electrodes for recording from the rat barrel cortex and images of the implanted µECoG electrode (left) and the µSEEG electrode (right). Note some contacts are outside brain tissue on the μSEEG electrode. Voltage responses across the left ribbon of the μECoG electrode to whisker stimulation at different whisker locations for the voltages and multi-unit activity (MUA). Green dots indicate significantly different from 0.5 sec before (which is baseline) air puff stimulation to the whisker (Wilcoxon rank-sum test) per channel and across trials. Number of trials >10.

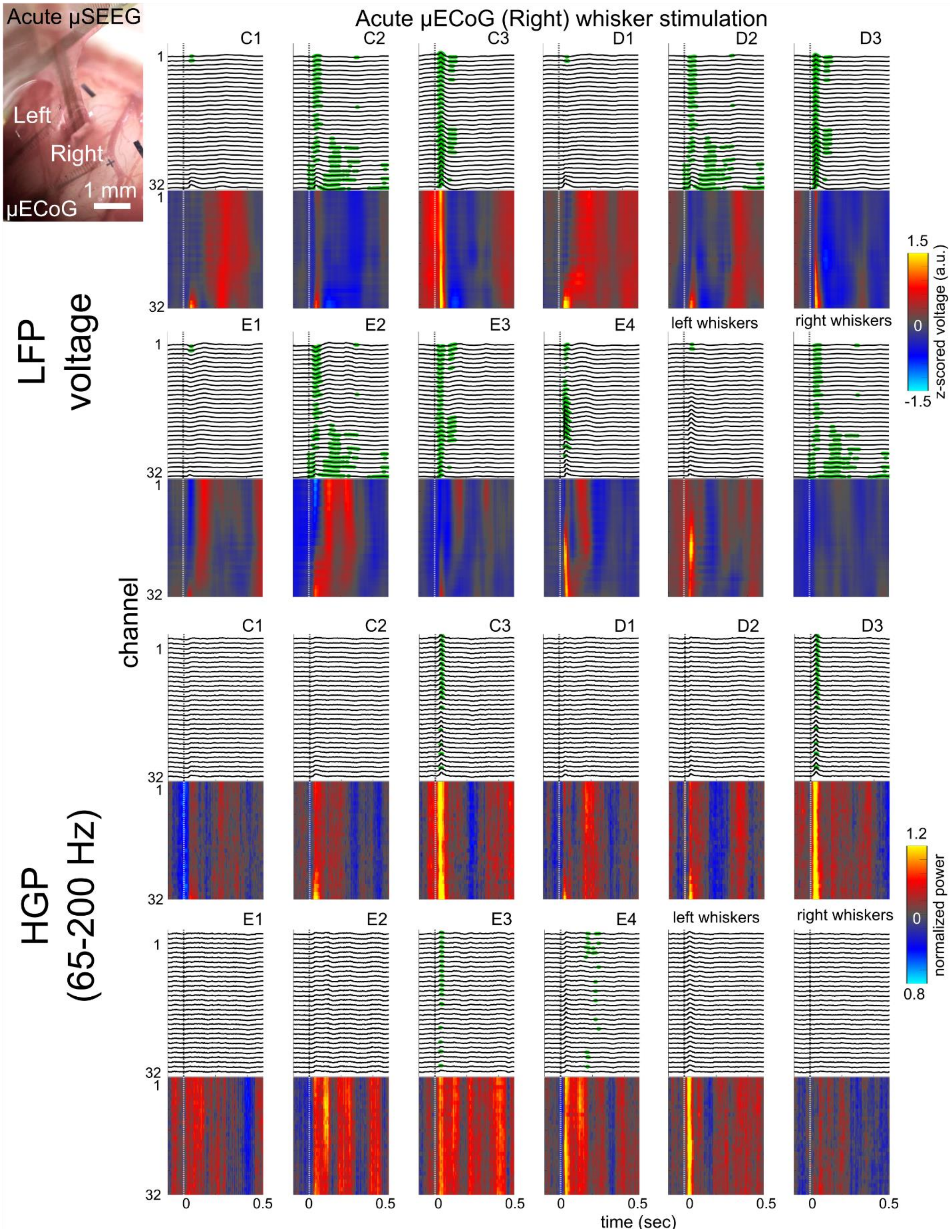


**Fig. S14. Acute surface recording with a strip of the μECoG over the rat barrel cortex with sensory stimulation.** Upper left: Location and 3D reconstruction of possible locations of the acute and chronic implantation of μSEEG electrodes for recording from the rat barrel cortex and images of the implanted μECoG electrode (left) and the μSEEG electrode (right). Note some contacts are outside brain tissue on the μSEEG electrode. Voltage responses across the right ribbon of the μECoG electrode to whisker stimulation at different whisker locations for the voltages and multi-unit activity (MUA). Green dots indicate significantly different from 0.5 sec before (which is baseline) air puff stimulation to the whisker (Wilcoxon rank-sum test) per channel and across trials. Number of trials >10.

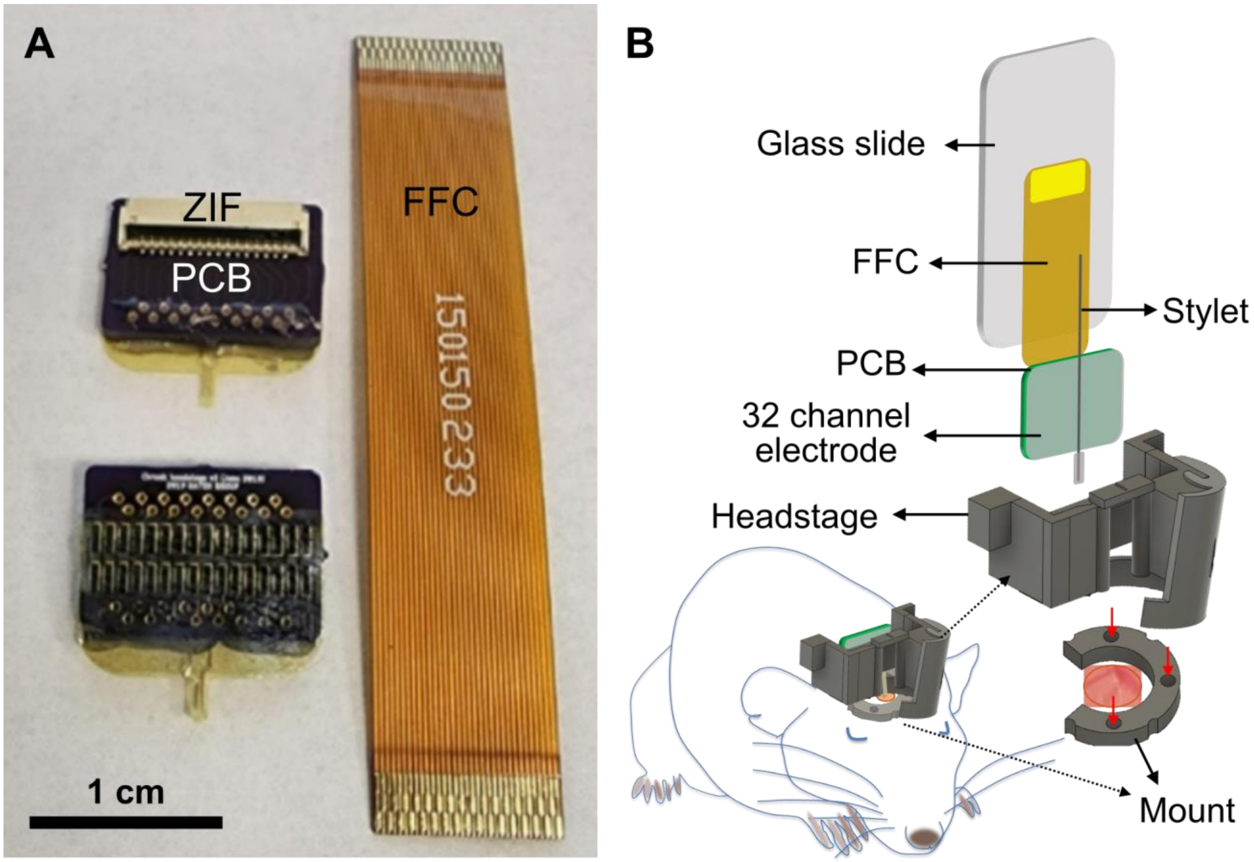


**Fig. S15.** Chronic implantation of the µSEEG electrode and recordings. (**A**) Hardware components and (**B**) Illustrations of the chronic µSEEG electrode implantation procedures.

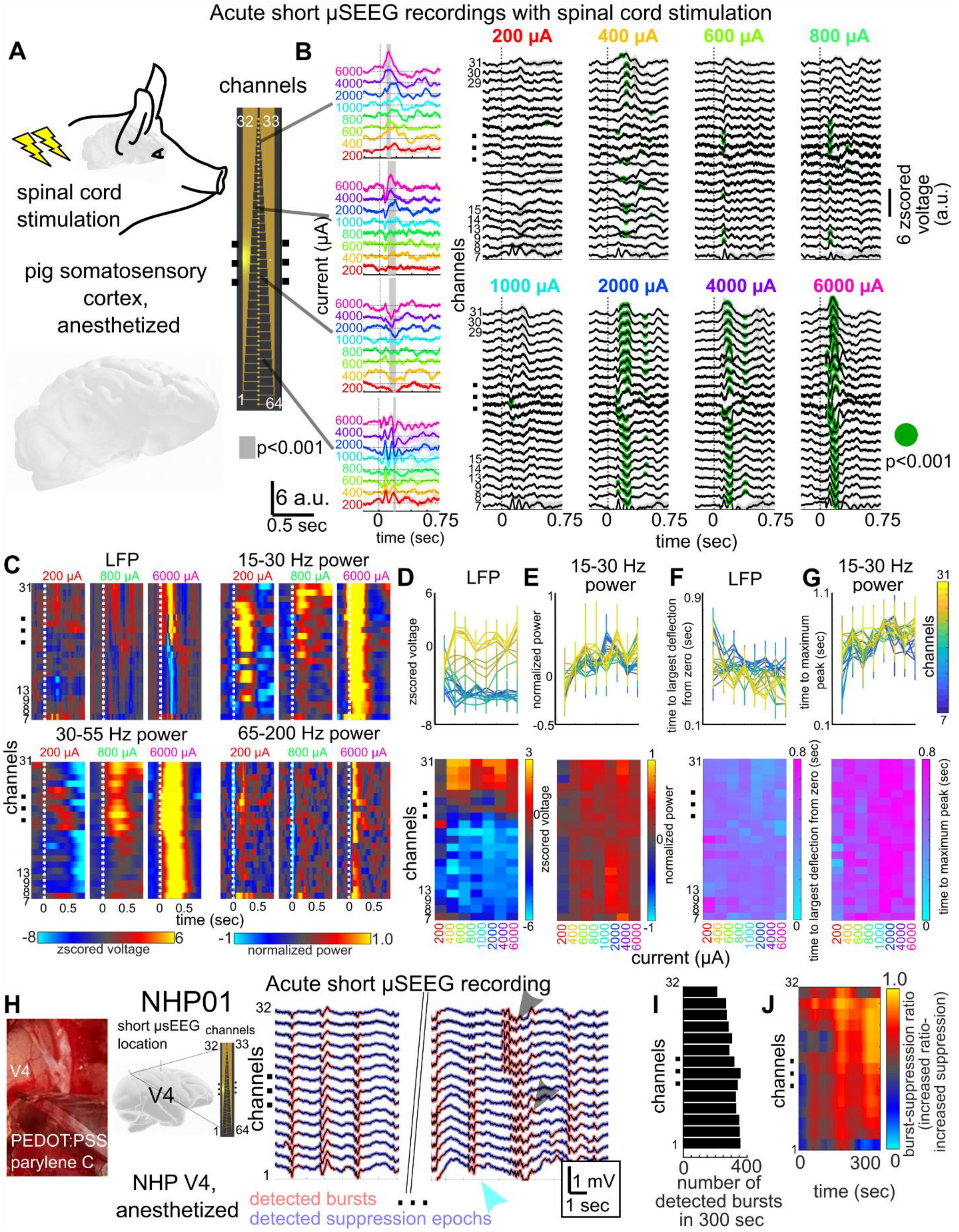


**Fig. S16. Acute short 64 channel µSEEG recordings in an anesthetized pig and non-human primate (NHP).** (**A**) Direct electrical stimulation of the spinal cord while doing an acute short 64 channel µSEEG electrode recording from the pig cortex, with responses increasing with increasing injected current. Grey bar indicates significantly different between current steps, Wilcoxon rank sum test. (**B**) Voltage responses along the electrode depth with more responses significantly different to baseline (0.5 sec before stimulation) occurring more with higher current levels (green dots, Wilcoxon rank sum test). (**C**) Average responses to stimulation in different frequency bands to different levels of injected current across channels. (**D**) Top: zscored voltage and Bottom: Two dimensional map of the time to zscored voltage per channel and per current step. (**E**) Top: 15-30 Hz power and Bottom: Two dimensional map of the time to 15-30 Hz power per channel and per current step. (**F**) Top: time to peak voltage and Bottom: Two dimensional map of the time to peak deflection from baseline per channel and per current step (**G**) Top: time to maximum deflection in 15-30 Hz power and Bottom: Two dimensional map of the time to maximum deflection in 15-30 Hz power per channel and per current step. (**H**) Acute implant of the short µSEEG electrode into V4 in an anesthetized NHP and ongoing evidence of burst suppression detected using automated approaches per channel. Arrowheads indicate detected bursts. (**I**) Number of detected bursts in 300 seconds per channel, with a correlation between channel depth and the number of bursts r= 0.72; p=0.0021 (Pearson's linear correlation). (**J**) burst-suppression ratio changes over time during the recording across channels.

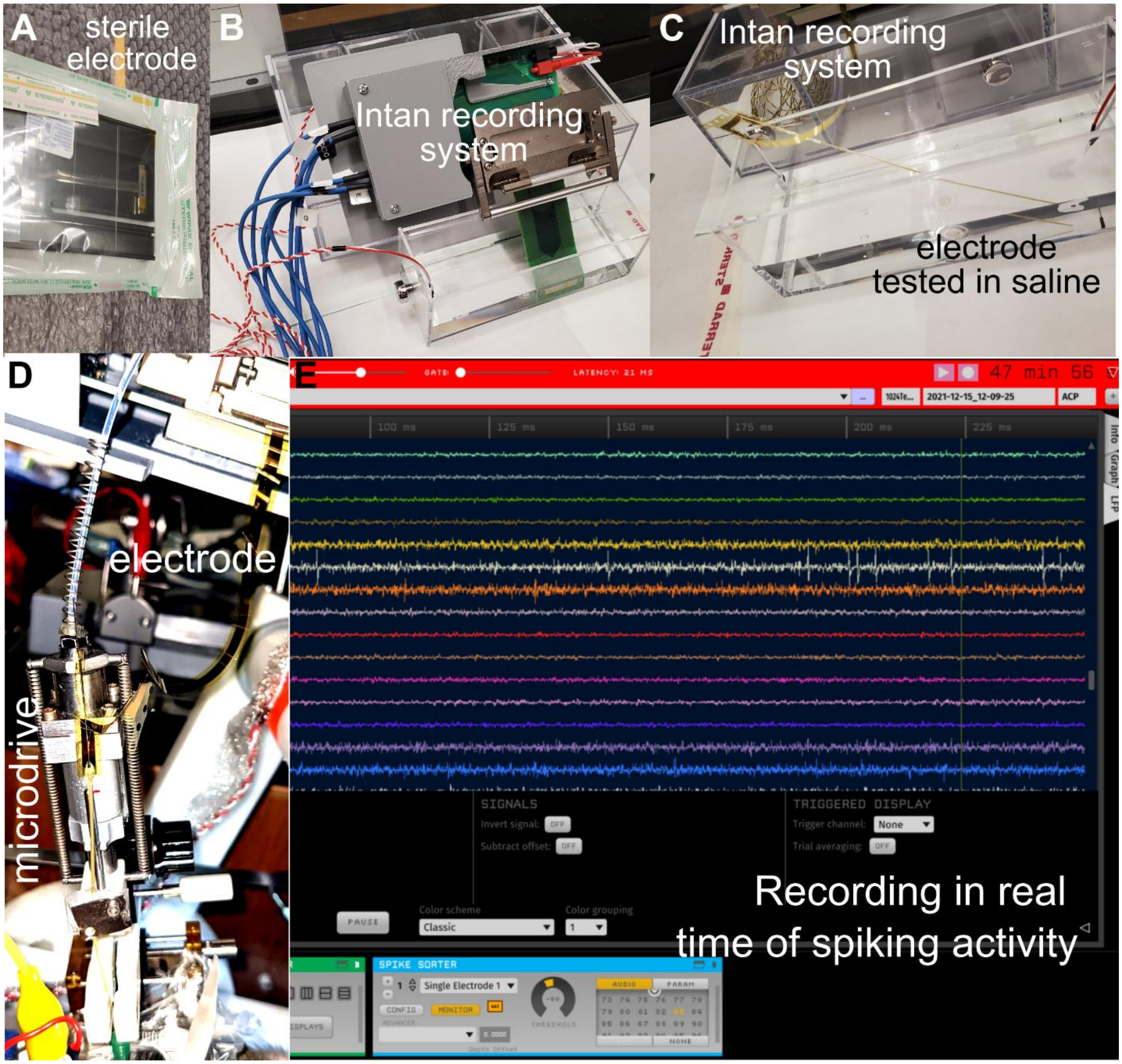


**Fig. S17. Set up and Microdrive recording using the long µSEEG electrode to record ongoing activity.** (**A**) Sterile electrode used in recording, sterilized using Ethylene Oxide gas. (**B**) 1024 channel Intan system used to record activity, testing in saline. (**C**) Testing of the long µSEEG electrode in saline. (**D**) Microdrive used to deliver the microelectrode. The same microdrive was used to drive tungsten electrodes to record activity. (**E**) Recording of real-time spiking activity using the long µSEEG electrode.

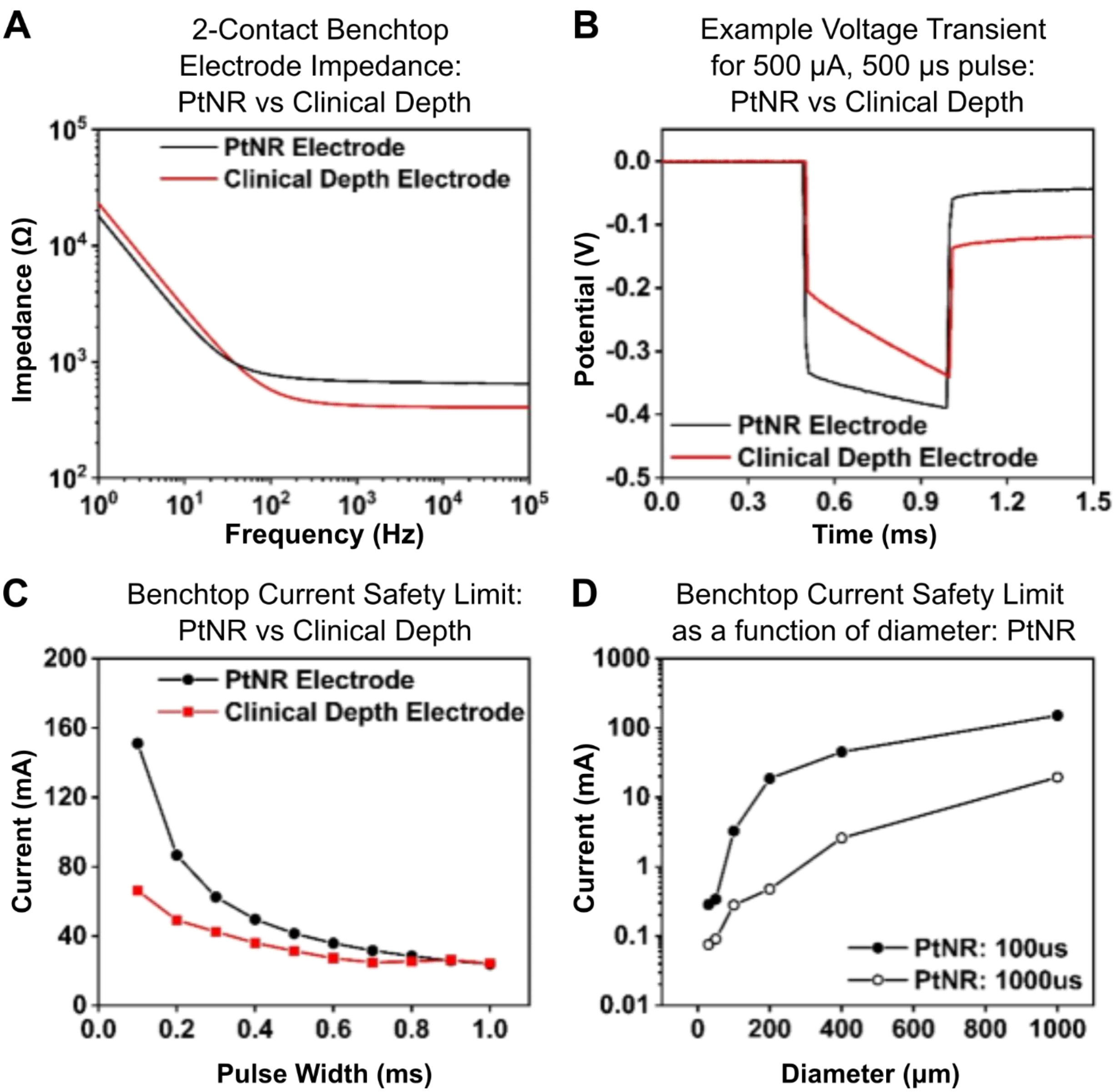


**Fig. S18. Electrochemical characterization of an AdTech Medical Spencer depth that is 1.12 mm diameter with contacts that are 2.41 mm long and 5 mm spaced and a PtNR electrode with 1 mm diameter during stimulation**. (**A**) Two-contact electrochemical impedance spectra (where another contact is used as both ground and reference to simulate impedances that appear in bipolar stimulation). The PtNR contact with 28.8 times smaller area than the PMT electrode exhibited higher capacitance and therefore lower reactance than the PMT electrode. The 1 kHz impedance that is typically overwhelmed with series resistances of metallization traces for macro-contacts is larger of the PtNR contact than the PMT electrode. (**B**) Example voltage transient across the two types of electrodes, exhibiting a lower potential drop across the electrode-tissue interface (non-instantaneous potential drop) for the PtNR contact compared to the PMT depth contact. (**C**) Benchtop electrochemical safety limits for the two types of electrodes when tested in the bipolar configuration (see ref. (*66*) for methods), showing superior performance for the PtNR contact, particularly at small pulse widths. (**D**) The benchtop electrochemical safety limits for two pulse widths for the PtNR contacts as a function of contact diameter.

**Table S1.** Electrode usage and species tested. Sp. – species, NHP- Non-Human primate, PEDOT-PEDOT:PSS, PI-Polyimide, PC-Parylene C.

| label | Sp. | Brain region | Electrode | PEDOT or PtNR | PI or PC | Acute or chronic |
|---|---|---|---|---|---|---|
| ***HS1*** | Human | Left anterior temporal lobe | Short 64 channel µSEEG | PEDOT | PC | Acute |
| ***HS2*** | Human | Left anterior temporal lobe | Short 64 channel µSEEG | PEDOT | PI | Acute |
| ***MM1*** | NHP | Visual cortex | Short 64 channel µSEEG | PEDOT | PC | Acute |
| ***MM1*** | NHP | Visual cortex | Surface µECoG | PtNR | PC | Acute |
| ***MM1*** | NHP | Visual cortex | Surface µECoG | PEDOT | PC | Acute |
| ***MM2*** | NHP | Parietal lobe/dlPFC and auditory cortex | Long µSEEG | PtNR | PI | Acute |
| ***MM2*** | NHP | Parietal lobe/dlPFC and auditory cortex | Tungsten electrode | - | - | Acute |
| ***RN1*** | Rat | Near barrel cortex | Short 32 channel µSEEG | PEDOT | PI | Chronic |
| ***RN2*** | Rat | Near barrel cortex | Short 32 channel µSEEG | PEDOT | PI | Chronic |
| ***RN3*** | Rat | Near barrel cortex | Short 32 channel µSEEG | PEDOT | PI | Chronic |
| ***RN4*** | Rat | Near barrel cortex | Short 32 channel µSEEG | PtNR | PI | Chronic |
| ***RN5*** | Rat | Near barrel cortex | Short 32 channel µSEEG | PtNR | PI | Chronic |
| ***RN6*** | Rat | Near barrel cortex | Short 32 channel µSEEG | PtNR | PI | Chronic |
| ***RN7*** | Rat | Near barrel cortex | Short 32 channel µSEEG | PtNR | PI | Chronic |
| ***RN8*** | Rat | Barrel cortex | Short 32 channel µSEEG | PtNR | PI | Chronic |
| ***RN9*** | Rat | Barrel cortex | Short 32 channel µSEEG | PtNR | PI | Chronic |
| ***RN10*** | Rat | Near barrel cortex | Short 64 channel µSEEG | PEDOT | PI | Acute |
| ***RN10*** | Rat | Near barrel cortex | Surface µECoG | PEDOT | PC | Acute |
| ***SS1*** | Pig | Brain | Short 64 channel µSEEG | PEDOT | PC | Acute |
| ***SS2*** | Pig | Brain | Short 64 channel µSEEG | PEDOT | PI | Acute |
| ***SS2*** | Pig | Brain | Surface µECoG | PEDOT | PC | Acute |
| ***SS3*** | Pig | Brain | Short 64 channel µSEEG | PEDOT | PI | Acute |
| ***SS3*** | Pig | Brain | sEEG electrode | - | - | Acute |
| ***SS3*** | Pig | Brain | Surface µECoG | PEDOT | PC | Acute |

**Table S2.** Electrode details.

| Electrode | Number of channels | Diameter (µm) | Center to center spacing (µm) | Recording length (mm) | PEDOT:PSS or PtNR | Polyimide (PI) or parylene C (PC) |
|---|---|---|---|---|---|---|
| **Short 32 channel µSEEG** | 32 | 30 | 60 | 1.89 | PEDOT:PSS, PtNR | PI |
| **Short 64 channel µSEEG** | 64 | 20 | 60 | 3.80 | PEDOT:PSS | PI, PC |
| **Long µSEEG** | 128 | 30 | 60 | 7.65 | PtNR | PI |
| **Surface µECoG** | 32 x 2 | 20 | 100 | 3.12 | PEDOT:PSS | PC |
